# Cell membrane proteome profiling uncovers molecular signatures and GPCR regulators underlying diet-induced obesity and exercise-mediated adaptation

**DOI:** 10.64898/2026.04.23.720492

**Authors:** Xiaoyu Lang, Lingyun Yang, Xiaoqi Lu, Qingqing Xu, Shanshan Li, Jie Yu, Huiting Luo, Liqin Guo, Xin He, Jiawen Liang, Hongbin Sun, Wei L. Shen, Wenqing Shui

## Abstract

Cell membrane proteins (CMPs), notably G protein-coupled receptors (GPCRs) and receptor tyrosine kinases (RTKs), represent the largest class of druggable targets. Despite their therapeutic importance, how the CMP landscape is systemically remodeled during metabolic disease progression and in response to exercise remains poorly understood. Here, we constructed the most comprehensive CMP proteome atlas to date across 22 male mouse tissues and established the first multi-tissue map of GPCR-associated protein hormones/prohormones. Integrated CMP proteomic and transcriptomic analysis revealed that diet-induced obesity and its progression to type 2 diabetes remodel the CMP repertoire in patterns distinct from those induced by exercise. We found that numerous GPCRs and RTKs critical for energy homeostasis are dysregulated during obesity in a tissue-specific manner, and these alterations are reversed to varying extents by exercise training. Importantly, our CMP-centric omics study identified two parathyroid hormone receptors (PTHRs) in the hypothalamus as negative regulators of feeding and body weight. Genetically silencing these hypothalamic PTHRs led to severe obesity even on a chow diet and markedly blunted exercise-induced weight loss. Collectively, our study provides not only new insights into systematic CMP regulation underlying metabolic deterioration *versus* exercise adaptation, but also nominates potential receptor targets for obesity management.

## Introduction

While obesity and type 2 diabetes (T2D) pose major global healthcare threat, regular exercise confers widespread health benefits, particularly by reducing risks of metabolic diseases, neurological diseases, and other pathologies^1, 2, 3^. In the last decade, various single- or multi-omics studies leveraging bulk or single-cell transcriptomics, epigenomics, proteomics and metabolomics have been conducted elegantly to interrogate the molecular mechanism underlying obesity/T2D development or exercise-induced adaptation^4, 5, 6, 7^. However, most previous studies are aimed to map the omic responses to either metabolic disease development or exercise training on healthy subjects, and often focus on a single or few tissues, with recent seminal work building molecular maps of multiple tissues^8, 9, 10^. Therefore, it remains largely unexplored regarding: (1) how obesity is differentiated from its associated severe morbidity T2D from a multi-tissue omics perspective, (2) how exercise training exerted during the progression of metabolic disorders systematically remodels the molecular architecture in diverse tissues.

Given that most biological processes are controlled in the protein dimension, unbiased and quantitative proteome profiling has been increasingly employed to gain unique insights into cellular function and disease mechanisms using animal models or human specimens^11, 12^. Within the proteome, cell membrane proteins (CMPs), which are integral components of the plasma membrane, are essential for sensing extracellular signals and maintaining cellular homeostasis, and they encompass several major protein families as therapeutic targets^13^. According to IUPHAR Guide to Pharmacology, these target-enriched CMP families primarily comprise G protein-coupled receptor (GPCR), catalytic receptor, ion channel, and transporter families. However, owing to their strong hydrophobicity, relatively low abundance and instability, CMPs are particularly challenging for conventional proteomics measurement, and are typically underrepresented in current MS-based proteome maps of human body or animal models^8, 9, 14, 15, 16, 17^. As a result, a high-coverage CMP proteome map is needed to expand the understanding of tissue-specific organization and disease-regulated dynamics of CMPs, which would facilitate the dissection of disease mechanisms and identification of therapeutic targets.

To address this key challenge, we previously developed an approach tailored to CMP proteome profiling in the region-resolved mouse brain by integrating several technical innovations^15^. Here, we extended this approach to build a comprehensive molecular atlas with the largest coverage of CMPs and their cognate protein/peptide ligands across 22 mouse tissues. We then profiled the CMP proteome and transcriptome dynamics across 12 tissues of mice during progression of obesity and T2D as well as exercise-induced adaptation. Our CMP omics study of both metabolic dysfunction and exercise adaption led to the discovery of two GPCRs in the hypothalamus, parathyroid hormone receptors PTH1R and PTH2R, as novel regulators of feeding behavior, diet-induced weight gain as well as exercise-mediated metabolic adaptation. While previous research has largely focused on the peripheral role of PTH1R in adiposity, we now uncover the metabolic functions of hypothalamic PTHR signaling. Furthermore, our study provides valuable resources for understanding the multi-tissue CMP proteome remodeling by obesity and T2D pathology in contrast to the proteome reversion by endurance exercise.

## Results

### A high-quality CMP proteome map of 22 male mouse tissues

To map CMP expression in various tissues, we collected 22 adult tissues of male C57BL/6J mice covering 12 major anatomical systems (Fig. 1a). Based on the previous protocol we established for cell membrane isolation, protein extraction and digestion for brain tissue^15^, we optimized these procedures in a tissue-specific manner to assure sufficient CMP recovery from each tissue prepared in three or four independent replicates (Supplementary Fig. 1). These CMP proteomic samples were then subjected to single-shot data-independent acquisition (DIA) MS analysis which gave rise to the most comprehensive CMP proteome map for mouse tissues (Fig. 1a). In total, 2,862 non-redundant protein groups classified as CMPs primarily from UniProt annotation (see Methods) were identified and quantified in at least one tissue, ranging from 1,348 to 2,151 CMPs profiled in different tissues (Fig. 1b; Table S1A). Our multi-tissue CMP atlas comprises a significant fraction (43.6%) of cell membrane-located therapeutic target family members including 311 GPCRs, 221 catalytic receptors, 224 ion channels, and 492 transporters (Fig. 1a). Compared to two recent large-scale global proteomic surveys profiled 41 mouse tissues^16, 17^, our proteome atlas excelled in the increased coverage of CMPs which reported 223 new CMPs and 89 new GPCRs not covered in previous mouse tissue proteome maps (Fig. 1c). Moreover, as a result of CMP enrichment and experimental optimization, superior quantification consistency for CMP and GPCR measurement across tissue replicates was achieved in our study (median coefficient of variation (CV), 9.4% for CMPs and 13.0% for GPCRs) compared to the previous study (median CV, 21.5% for CMPs and 26.7% for GPCRs) (Fig. 1d and Supplementary Fig. 2a).

**Figure 1.**
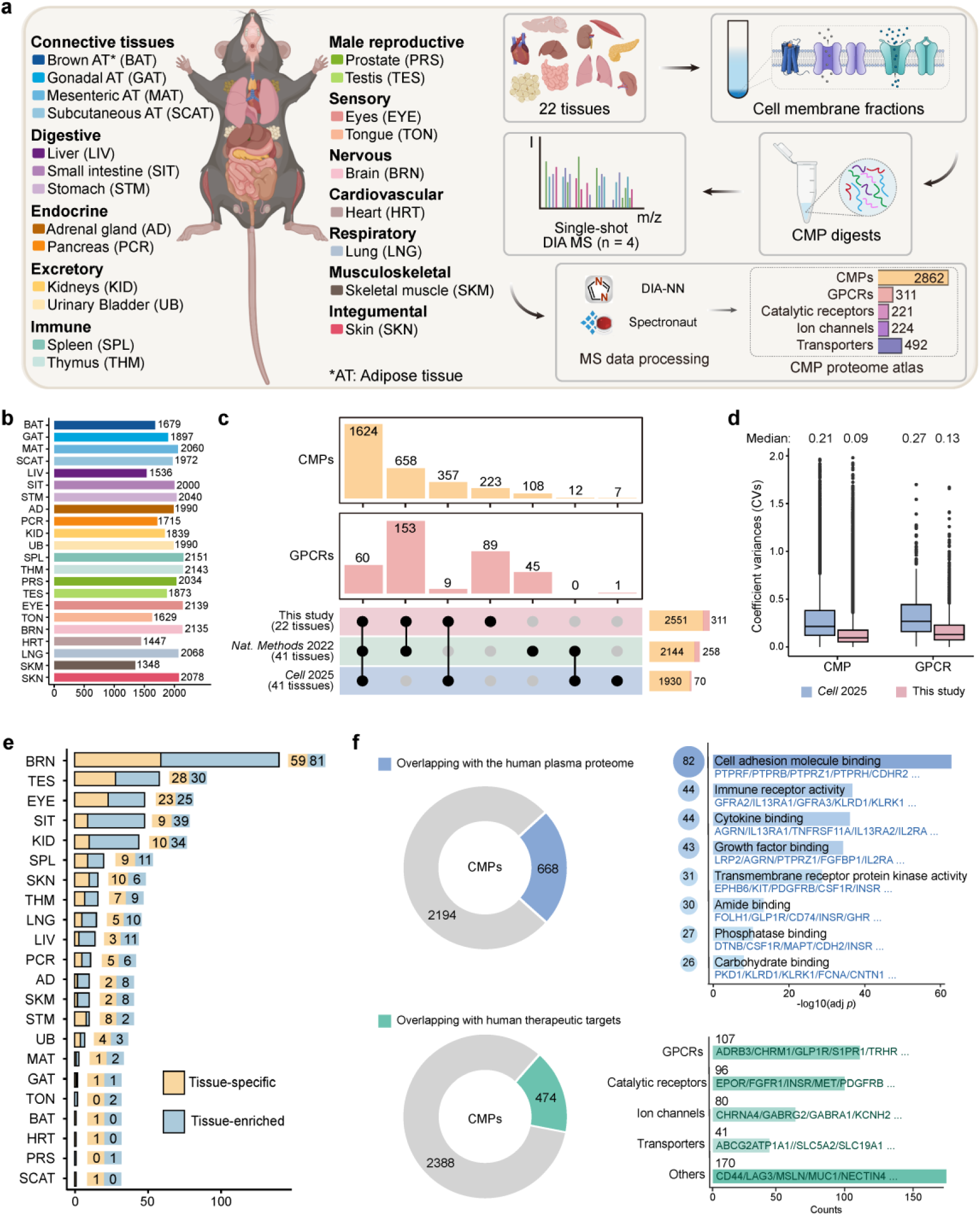
Generating a CMP proteome map across male mouse tissues. (a) Schematic workflow for cell membrane protein (CMP) proteome profiling of 22 male mouse tissues and the total number of CMP, GPCR, catalytic receptor, ion channel and transporter identifications in the map. (b) Number of CMPs quantified in different tissues. (c) Overlapping and unique CMPs and GPCRs profiled in this study and those in previous mouse tissue proteome maps^16, 17^. The stacked bars on the right indicate the number of GPCR (pink) and other CMPs (organge) reported in each dataset. (d) Reproducibility of CMP and GPCR protein quantification in this study *vs* the previous study by Lu T *et al*^17^. as represented by coefficient variances (CVs) of protein intensity between replicates for all tissues. (e) Number of tissue-specific (only detected in one tissue) and tissue-enriched (> 5-fold higher expression than any other tissues) CMPs in each tissue. (f) Overlap between our multi-tissue CMP proteome and the human plasma proteome^18^ (upper) or between our CMP proteome and curated therapeutic targets^19^. Enriched GO-MF terms of the overlapping proteins or major categories of CMP drug targets are shown on the right.

Our comprehensive and quantitative CMP map allowed us to uncover 189 CMPs that are specifically expressed in one tissue (not detected in any other tissues) and 289 CMPs that are significantly enriched in one tissue (> 5-fold higher expression than any other tissues). Top three tissues containing the most tissue-specific and tissue-enriched CMPs are brain, testis, and eye (Fig. 1e). Notably, these tissue-specific/enriched CMPs are primarily enriched in biological processes that are reflective of the fundamental and unique functions of specific organs (Supplementary Fig. 2b). In addition, by overlapping our multi-tissue CMP proteome map with the human plasma proteome^18^ and the therapeutic target database^19^ for proteins evolutionally conserved between mice and humans, we are able to infer the tissue origin of 668 CMPs present in human plasma and 474 CMPs serving as targets for approved or clinical trial drugs (Fig. 1f). Our analysis indicates an average of 26.7% CMPs expressed in individual tissues are possibly released into plasma and an average of 16.5% tissue-expressed CMPs act as therapeutic targets for various diseases (Supplementary Fig. 2c).

### Transcriptome and proteome maps for CMPs

To gain insights into CMP abundance regulation at the transcript and protein levels, we also profiled the transcriptomes for the exact same tissues collected for proteomic analysis, which led to the quantification of 3,237 CMP-corresponding transcripts expressed in at least one tissue (Table S1B). Both the principal components analysis (PCA) and the hierarchical clustering tree based on our CMP proteome and transcriptome profiling revealed differential patterns of tissue clustering or segregation (Fig. 2a and Supplementary Fig. 3a). While tissues in the proteome map are more evenly distributed and form several sub-clusters, the majority of tissues in the transcriptome map are clustered more tightly with only a few such as brain and eye deviated away (Fig. 2a). Furthermore, the tissue correlation analysis suggests that brain, eye, and testis are top 3 tissues that exhibit the most distinct CMP abundance patterns compared to other issues, with more variability observed at the mRNA level than the protein level (Fig. 2b). Within individual tissues, an overall modest correlation between mRNA and protein abundance was observed for all CMPs (Spearman correlation *r* = 0.42-0.58) whereas tissue-specific or enriched CMPs exhibited higher variability in the mRNA-to-protein correlation (*r* = 0.11-0.68) (Fig. 2c). Additionally, among 1,922 CMPs co-expressing their mRNA and protein in at least five tissues, 14.2% (273 CMPs) showed weak across-tissue mRNA-to-protein correlation (*r* < 0.3) and their molecular functions are enriched in metal ion transmembrane transporter activity, phospholipid binding, actin binding, cell adhesion molecule binding, etc. (Supplementary Fig. 3b, c).

**Figure 2.**
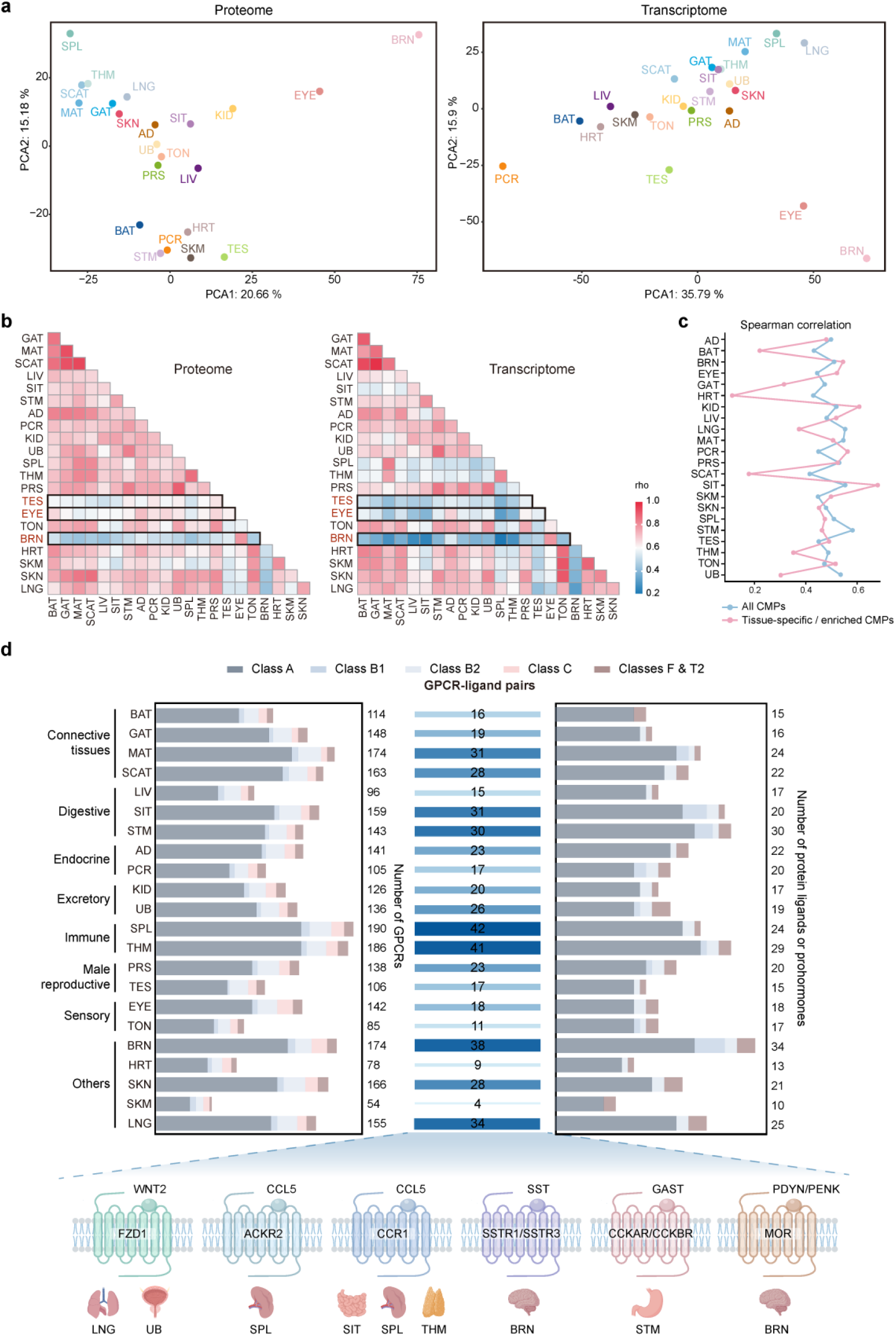
Multi-tissue distribution of CMP proteome, transcriptome, and GPCRs in association with their cognate protein or peptide ligands. (a) PCA showing differential patterns of tissue clustering or segregation based on the CMP proteome data (left) *vs* the CMP transcriptome data (right). (b) Spearman correlation of CMP molecular abundances at the proteome (left) or transcriptome (right) levels across 22 tissues. Three tissues with the lowest correlation to the other tissues are highlighted in black boxes. (c) Spearman correlation between proteomic and transcriptomic measurements for all CMPs (blue line) or tissue-specific/enriched CMPs (pink line) within each tissue. (d) Number of GPCRs in different classes (left) and the number of protein ligands or prohormones associated with these GPCRs (right) profiled in different tissues, and the number of GPCR-ligand pairs detected in different tissues (middle). The total numbers covering all GPCR classes are annotated beside each bar. Representative pairs of GPCRs and cognate protein ligands that are co-identified in specific tissues are illustrated at the bottom.

Taken together, the comparison of CMP transcriptome and proteome maps, aligning with previous global multi-omics profiling of animal and human tissues, points out the contribution of factors other than mRNA levels to controlling cell membrane protein abundance and homeostasis in various tissues^14, 16, 20, 21^.

### Multi-tissue profiling of GPCRs and their endogenous protein/peptide ligands

As the largest cell membrane protein family encoded in the human genome, GPCRs mediate a wide range of signaling processes and are targeted by one third of the drugs in clinical use^22, 23^. For GPCRs recognizing protein or peptide hormones as their endogenous ligands, some of them with disease implications are successfully targeted by peptide therapeutics (*e.g.* μ and κ opioid receptors in pain relief, apelin and angiotensin receptors in cardiovascular disease, and GLP-1 receptor in obesity and type 2 diabetes)^22, 24, 25, 26^. Given that the membrane fraction was extensively washed in our experiment to retain integral membrane proteins and proteins strongly associated with the membrane, we reasoned that our CMP proteomic analysis enables not only profiling of membrane receptors such as GPCRs but also capturing their endogenous protein or peptide ligands.

Focusing on the GPCR system, we first analyzed the tissue distribution of all measured non-redundant GPCR proteins that fall into six classes according to their structural similarity (Fig. 2d)^27^. Two immunity-related tissues, thymus and spleen, possess the highest number of GPCR proteins (186 and 190), followed by mesenteric adipose tissue and brain. On the other side, for protein and peptide receptors mostly belonging to classes A, B1 and F, our data analysis reveals the multi-tissue profile of 49 cognate protein ligands for these receptors and 12 prohormones that undergo specific proteolysis to yield peptide ligands associated with these receptors (Table S2). Brain, stomach and small intestine are top 3 tissues having the most identifications of cognate protein ligands or prohormones for GPCRs. Presumably, these protein or peptide ligands are either produced within the tissue or elsewhere before trafficking to the tissue. To infer potential activation of GPCRs upon binding to their endogenous agonists, we analyzed paired GPCRs and their protein/peptide ligands that are both detected in the same tissue in our CMP proteome map. These GPCR-ligand pairs retained in the tissue-derived membrane fractions indicate the protein or peptide hormones may be bound to their cognate receptors on the cell membrane to mediate native GPCR activities. For example, protein ligand WNT2 was co-identified with its receptor Frizzle 1 (FZD1) in the lung and urinary bladder (Fig. 2d). Chemokine CCL5 was co-identified with one receptor ACKR2 exclusively in spleen yet with another receptor CCR1 in three tissues (small intestine, spleen, and thymus). Somatostatin (SST) and gastrin (GAST), two protein ligands known to act on multiple receptors, were co-identified with receptors SSTR1/SSTR3 in the brain and with receptors CCKAR/CCKBR in the stomach, respectively. Our study also identified prohormones proenkephalin-A (PENK) and proenkephalin-B (PDYN) which yield neuropeptides that can activate opioid receptors to be co-expressed with the μ-opioid receptor (MOR) in the brain (Fig. 2d).

Collectively, our result suggests that 4 to 42 GPCRs recognizing protein or peptide hormones may be associated with and potentially activated by their cognate ligands present in individual tissues (Fig. 2d). These GPCRs constitute 18%–65% of all peptide/protein receptors detected in different tissues (Supplementary Fig. 3d). Therefore, our study generates a multi-tissue co-regulatory map of membrane receptors and their protein/peptide ligands for the GPCR superfamily.

### Multi-tissue CMP proteomic responses to obesity, diabetes and exercise training

To provide new insights into the molecular mechanisms underpinning chronic metabolic diseases, we employed our CMP proteome profiling approach to characterize the multi-tissue proteomic responses to obesity and T2D using high-fat diet (HFD)-induced mouse models. Meanwhile, to understand the wide-ranging health benefits provided by regular exercise to alleviate metabolic disease symptoms^1, 9, 10^, we also profiled CMP proteomic responses of mice subjected to endurance exercise training. To this end, we developed mouse models in four groups, all starting with 10-week-old mice (Fig. 3a): (1) in the chow group, mice were fed with chow diet for 9 weeks; (2) in the obesity group, mice were fed with high-fat diet (HFD) for 9 weeks; (3) in the exercise group, mice fed with HFD were subjected to treadmill endurance exercise training for 6 weeks; (4) in the T2D group, mice were fed with HFD for 13 consecutive weeks. At the study endpoint, the obesity group significantly increased body weight by 38.3% and blood glucose by 18.4% compared to the chow group (Fig. 3b). The T2D group further increased the body weight by 20.9% and the blood glucose level by 15.6% relative to the obesity group, and showed impaired glucose tolerance (Fig. 3b and Supplementary Fig. 4a). The beneficial effect of exercise training was demonstrated by the significant reduction of body weight by 14.5% and blood glucose by 14.1% for the exercise group compared to the obesity group (Figures 3b and S4B). We then dissected 12 metabolically relevant tissues (hypothalamus, liver, small intestine, pancreas, skeletal muscle, brown adipose tissue (BAT), gonadal adipose tissue (GAT), mesenteric adipose tissue (MAT), subcutaneous adipose tissue (SCAT), spleen, kidney, heart) from mice in four groups in biological triplicate, and isolated their cell membrane fractions for proteomic analysis. At the meantime, these tissues were separately collected from three groups (chow, obesity and exercise) for bulk transcriptomic analysis. Notably, unlike the previous 22-tissue CMP proteome map covering the whole brain, here we collected the hypothalamus which is a critical brain region for regulating energy homeostasis^28^. From SDS-PAGE images of CMP extracts, most tissues displayed very similar protein patterns among four groups except for liver, of which the T2D group apparently differs from other three groups (Fig. 3c and Supplementary Fig. 4c). After proteolysis, these CMP proteomic samples were subjected to both single-shot DIA MS analysis and peptide pre-fractionation prior to DDA MS analysis for different tissues. By merging DIA and DDA data from all tissues under all conditions (240 MS raw files), we constructed the most comprehensive hybrid spectral libraries for CMPs in the disease and exercise models. DIA data were then processed using two complementary search engines (DIA-NN and Spectronaut) with the hybrid libraries, which resulted in the identification and quantification of 2698 CMPs in at least one tissue for one group, with excellent quantification consistency between tissue replicates for all groups (median CV 7.23-7.57%) (Supplementary Fig.5a).

**Figure 3.**
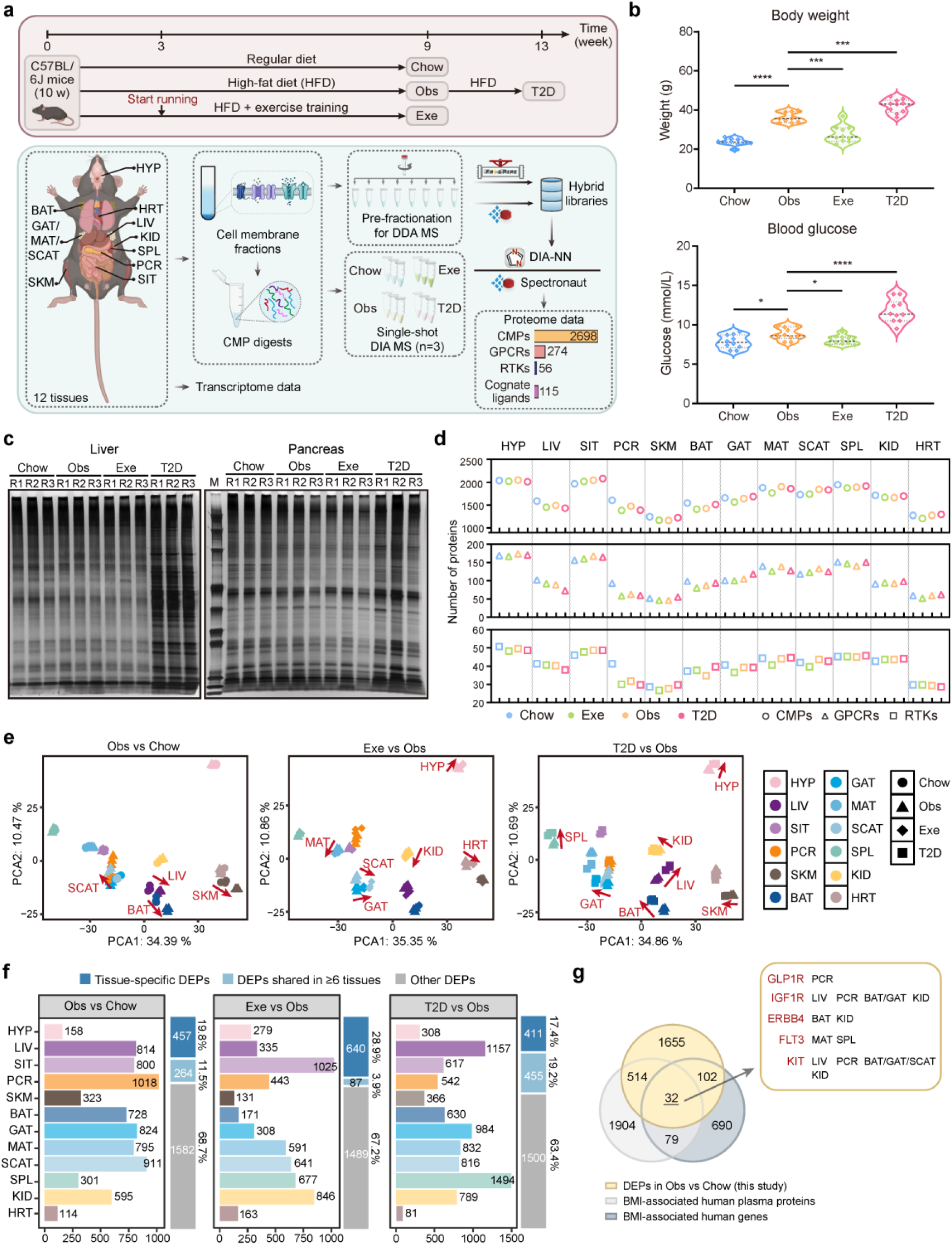
Multi-tissue CMP proteome profiling in mouse models of obesity, diabetes, and exercise training. (a) Overall workflows illustrating the establishment of mouse groups of obesity (Obs), diabetes (T2D), exercise (Exe) and control (Chow) (upper) and CMP proteome/transcriptome profiling experiments across 12 tissues of mouse models (lower). The total number of CMPs, GPCRs, RTKs, and their associated ligands present in the proteome data are shown in the bar graph. (b) Measurement of body weight and fasting blood glucose for the Chow group (n = 10, blue), Obs group (n = 9, orange), Exe group (n = 10, green), and T2D group (n = 9, pink) at the end of model development. Data were analyzed by unpaired two-tailed t-tests between Obs vs Chow, Exe vs Obs and T2D vs Obs groups. \**p* < 0.05, \*\*\**p* < 0.001, \*\*\*\**p* < 0.0001. (c) SDS-PAGE images of CMP total extracts prepared from liver (left) or pancreas (right) from four experimental groups in three independent replicates. (d) The average numbers of CMP, GPCR and RTK protein identifications in different tissues from four experimental groups. (e) PCA showing differential patterns of tissue molecular architecture in the three pairwise comparisons based on CMP proteome data from four groups. Red arrows denote a significant separation of pair-wise groups for specific tissues (*p* < 0.01) responding to the obesity development (left), exercise training (middle), or obesity-to-T2D progression (right). (f) Number of differentially expressed CMPs (DEPs) identified in each tissue across three pairwise comparisons, highlighting the number and proportion of tissue-specific or multi-tissue shared DEPs for each comparison. (g) Venn diagram depicting overlaps between DEPs from all tissues regulated in the obesity development (this study) and two lists of BMI-associated human plasma proteins^18^ and BMI-associated human genes^36, 37^. Five GPCRs or RTKs in the central overlap of three lists with their tissues of regulation in the Obs vs Chow comparison are specified.

Importantly, this dataset encompasses proteomic responses of 274 GPCRs and 56 receptor tyrosine kinases (RTKs), two membrane receptor families known to play pivotal roles in energy homeostasis with quite a few members emerging as anti-obesity or anti-diabetic drug targets (Fig. 3a, d) ^29, 30, 31, 32, 33^. In addition, by searching all known protein hormones or prohormones associated with these GPCRs or RTKs, our study profiled the abundance of 115 GPCR or RTK ligands in different tissues of the metabolic disease or exercise models (Fig. 3a).

### Differentially expressed CMPs responding to disease progression or exercise training

To understand CMP proteome remodeling by disease progression and reversion by exercise, we focused our data mining on three pair-wise comparisons, obesity vs chow, exercise vs obesity, and T2D vs obesity. The comparisons of obesity vs chow and T2D vs obesity would reveal CMP proteomic perturbation occurring in the development of obesity and in the progression from obesity to T2D, respectively. On the other hand, the comparison of exercise vs obesity would highlight proteomic adaptation to exercise training that may reduce risks of obesity. PCA of these pair-wise experimental groups based on all CMP proteome data revealed clustering of all replicates from paired groups for most tissues, suggesting the tissue-specific CMP molecular architecture was mainly retained in the disease and exercise groups relative to the chow group (Fig. 3e). However, we also observed separation of pair-wise groups for certain tissues (*P* < 0.01) as annotated by red arrows in Figure 3e. For example, liver, skeletal muscle, and BAT shifted their CMP molecular signatures both from chow to obesity and from obesity to T2D groups, supporting the notion that metabolic functions of these tissues are dysregulated in obesity and likely exacerbated in T2D^2, 34, 35^. Additionally, exercise training induced the separation of specific tissues including hypothalamus, heart, and multiple adipose tissues between obesity and exercise groups, indicating the distinct effects of endurance exercise in remodeling tissue CMP proteomes (Fig. 3e).

Next, for three pair-wise comparisons, we identified sets of differentially expressed CMPs (DEPs) which are classified into tissue-specific DEPs or multi-tissue shared DEPs (Fig. 3f; Table S3). In addition, we calculated percentages of DEPs over all measured CMPs in individual tissues (Supplementary Fig. 5b). Interestingly, the median DEP percentages across all tissues are 35.5% in obesity vs chow, 21.1% in exercise vs obesity, and 35.9% in T2D vs obesity comparisons, which suggests that extensive multi-tissue CMP regulation occurs in the progression of both obesity and T2D whereas exercise training remodels CMP proteome responses to a less extent (Supplementary Fig. 5c). We then compared tissue-specific sets of DEPs with those of differentially expressed CMP genes (DEGs) identified from transcriptomics. Surprisingly, the fraction of DEPs overlapping with DEGs in the same direction of regulation and in the same tissue was rather low, in the range of 0%-26.8% for different tissues in the obesity vs chow comparison and 0%-24.9% in the exercise vs obesity comparison (Supplementary Fig. 5d). Moreover, protein-to-mRNA correlation of CMP abundances varied from 0.42 to 0.56 for different tissues at different conditions (Supplementary Fig. 5e). Thus, the marked discordance of DEP and DEG selection from our proteomic and transcriptomic data, which was also reported in several multi-omics studies^8, 9, 10^, highlights the prevalent post-transcriptional regulation of CMPs that may be critical to establishing tissue-specific proteostasis in response to pathophysiological perturbation.

To evaluate the translational value of our data, we collected lists of 903 human genes associated with body mass index (BMI) from two GWAS datasets^36, 37^ and 2,529 human plasma proteins also associated with BMI from a plasma proteome dataset^18^. By comparing these two lists with a total of 2,303 DEPs we identified from mouse tissues regulated in the obesity development, we observed a significant fraction of our obesity-induced DEPs (28.1%) overlapping with BMI-associated human genes or BMI-associated human plasma proteins (Fig. 3g). Furthermore, within the overlap of 32 homologous proteins/genes shared by these three lists, we observed GLP1R, a prime GPCR target for obesity and T2D therapeutics^31^, as well as two RTKs (IGF1R, ERBB4) known to regulate glucose uptake, insulin sensitivity or adipose inflammation, and contribute to obesity-related metabolic dysfunction^38, 39^. These results jointly support an appreciable concordance of our data from mouse models with human studies and their relevance to human metabolic disorders.

To reveal tissue-specific regulation of CMPs with central roles in metabolic homeostasis, we examined the abundance changes of several membrane receptors and a panel of glucose transporters in the three pairwise comparisons (Fig. 4a). For leptin receptor detected in the MAT and insulin receptor in liver, skeletal muscle, and SCAT, they were all down-regulated in obesity vs chow and up-regulated in exercise vs obesity comparisons, indicating their obesity-induced dysfunction could be in part ameliorated by exercise training. Conversely, for the protein hormones stimulating these receptors, leptin detected in all four adipose tissues and insulin in pancreas, largely increased their levels in the development of obesity while reduced levels in the respective tissues responding to exercise. Moreover, leptin levels were further increased in BAT and MAT in the obesity-to-T2D progression while its levels in GAT and SCAT and insulin in pancreas all remained unchanged in this process. These findings not only align well with the fact that both leptin resistance and insulin resistance contribute to overeating, weight gain and hyperglycemia in obese or T2D animals and humans^40, 41^, but also reveal molecular adaptation of multiple tissues to endurance exercise. To highlight those CMPs or their ligands that are regulated in one direction in the obesity vs chow comparison and in the opposite direction in the exercise vs obesity comparison, we refer to them as reverse-trend proteins which are annotated with arrows in opposite directions in the CMP regulation heat map (Fig. 4a). Interestingly, GLP-1 receptor which is highly expressed in pancreatic β cells and acts as a key regulator of insulin secretion and glucose homeostasis^31, 42^, is not a reverse-trend protein as it was down-regulated in pancreas in the obesity development yet not reversed by exercise training. This result implies obesity-related β cell dysfunction cannot be fully rescued with the exercise training protocol used in this study.

**Figure 4.**
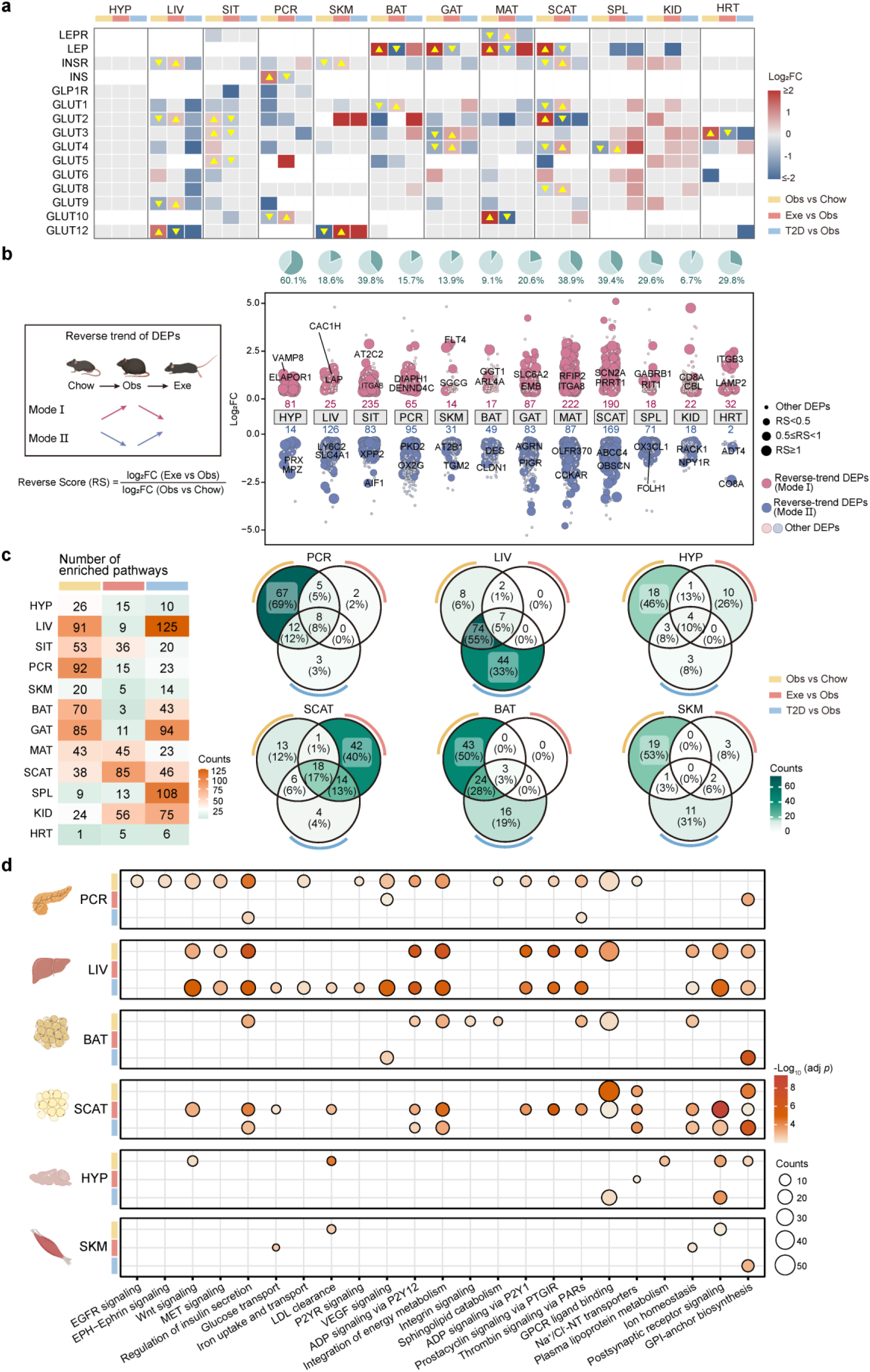
Reverse-trend CMPs and tissue-specific pathways enriched in obesity, diabetes, or exercise interventions. (a) Heatmap illustrating the regulation of three obesity-related membrane receptors, and two associated protein hormones, and ten glucose transporters in different tissues in the comparisons of Obs vs Chow, Exe vs Obs, and T2D vs Obs. Color code denotes the log_2_FC range and the trend of regulation (red, up-regulated; blue, down-regulated; light gray, detected yet not significantly regulated). Pairs of yellow triangles denote reverse-trend proteins with opposite directions of regulation for two pairwise comparisons. FC, fold change. (b) Illustration of two modes of reverse-trend and the calculation of reverse score (RS) (left), and distribution of reverse-trend CMPs across different tissues (right). Reverse-trend CMPs are highlighted with dark colored bubbles (bubble size indicating the RS and the numbers shown next to the tissue name). CMPs with top2 RS in each mode are specified for each tissue. The pie charts above depict the percentages of reverse-trend CMPs among all obesity-regulated CMPs in each tissue. (c) Number of enriched REACTOME pathways (adj. *p* < 0.01) based on DEPs identified from each tissue in three pairwise comparisons or three interventions (left). Venn diagrams showing unique and overlapping pathways specifically enriched in different interventions for six tissues (right). (d) Representative CMP-derived pathways specifically enriched (adj. *p* < 0.01) in the six tissues responding to three interventions.

In addition, our study uncovers disease or exercise-regulated patterns of 10 glucose transporters in various peripheral tissues (Fig. 4a). Among them, GLUT4 in skeletal muscle from T2D patients or other metabolic disorders is known to reduce its translocation to the plasma membrane as a result of impaired glucose uptake^43, 44^. Interestingly, our study revealed a decrease of the cell membrane abundance of GLUT4 in seven tissues (*i.e.* liver, pancreas, skeletal muscle, spleen, heart and two adipose tissues) in the obesity vs chow comparison, suggesting the attenuated translocation of GLUT4 could happen in a wide range of tissues. Additionally, GLUT4 showed a reverse-trend of regulation in the exercise vs obesity comparison in three tissues (GAT, SCAT and spleen). For other GLUT proteins, they were also found to be reverse-trend proteins in selected tissues responding to obesity development and exercise training. Moreover, five GLUT proteins (GLUT1/2/4/5/9) in specific tissues were responsive to obesity and T2D progression in the same direction of regulation.

We then systematically analyzed the distribution of reverse-trend CMPs across different tissues. Depending on the direction of regulation, these CMPs exhibited reverse trends in mode I or mode II, with the extent of exercise-induced reversing effects indicated by a reverse score (Fig. 4b left). SCAT is the tissue having the largest number of reverse-trend CMPs, whereas heart has the lowest number (Fig. 4b right). Notably, the percentage of reverse-trend CMPs over all obesity-regulated CMPs in each tissue varies from 6.7% in kidney to 60.1% in hypothalamus with a median of 26.9% (Fig. 4b right). Therefore, we conclude that exercise training could widely yet partially reverse the multi-tissue CMP proteome landscape dysregulated by obesity whereas T2D pathogenesis may exacerbate the dysregulation of specific CMPs (*e.g.*, leptin receptor, insulin receptor, and multiple glucose transporters) in selected tissues.

### Tissue-specific pathways enriched in disease or exercise interventions

Pathway enrichment based on DEPs identified from each tissue in three pairwise comparisons (referred to as three interventions) gave rise to differential sets of pathways enriched in individual tissues during the interventions of obesity, T2D or exercise training (Fig. 4c left). Pancreas and liver have the largest number of CMP-derived pathways enriched in obesity and T2D interventions, respectively (92 for pancreas in the obesity vs chow comparison, 125 for liver in the T2D vs obesity comparison) while SCAT is the top1 tissue with the most pathways enriched in the exercise intervention (85 in the exercise vs obesity comparison).

When comparing the sets of pathways specifically regulated in any of the three interventions for a given tissue, pancreas has a much larger proportion of unique pathways exclusively enriched in the obesity intervention (69%) than unique pathways enriched in exercise (2%) and T2D (3%) interventions (Fig. 4c right). These obesity-enriched unique pathways in pancreas include Wnt/β-catenin signaling, EPH-Ephrin signaling, EGFR signaling, MET signaling, and insulin secretion (Fig. 4d, Table S4), suggesting obesity specifically induces wide perturbation of membrane receptor-mediated diverse signaling that are critical for β-cell survival, proliferation, and insulin secretion^45, 46, 47, 48^. Given these pathways were not enriched in exercise and T2D interventions, it indicates they were not significantly rewired in pancreas by exercise training or during obesity-to-T2D progression. In contrast, liver responded most extensively to the T2D intervention with 33% unique pathways enriched in this intervention compared to those enriched in obesity (6%) and exercise (0%) interventions (Fig. 4c right). These T2D-enriched unique pathways involve P2YR signaling, Wnt signaling, VEGF signaling, glucose transport, iron uptake and transport, and LDL clearance (Fig. 4d, Table S4), suggesting obesity-to-T2D progression widely rewire membrane receptor-coupled signaling to affect glucose and lipid metabolism in liver^49, 50, 51, 52, 53, 54^.

In addition, when comparing BAT and one type of white adipose tissue SCAT, they exhibited distinct patterns of CMP-derived pathway enrichment. BAT responded most extensively to the obesity intervention with 50% unique pathways enriched in this intervention, yet no unique pathway was enriched in the exercise intervention (Fig. 4c right). Obesity-enriched unique pathways in BAT include integrin signaling, ADP signaling via P2Y12, regulation of insulin secretion, sphingolipid catabolism, and integration of energy metabolism (Fig. 4d, Table S4), most of which contribute to dysfunction of glucose/lipid metabolism or thermogenesis^55, 56, 57, 58, 59^. SCAT, however, responded modestly to the obesity intervention with 12% unique pathways enriched, while exercise induced a much stronger response with 40% unique pathways enriched in this intervention (Fig. 4c right). These exercise-enriched unique pathways in SCAT involve prostacyclin signaling via PTGIR, thrombin signaling via PARs, ADP signaling via P2Y1, glucose transport, and LDL clearance (Fig. 4d, Table S4), which are known to mediate lipolytic, anti-inflammatory or insulin-sensitizing effects^60, 61, 62, 63^. The rest of tissues also display specific patterns of pathway enrichment in different interventions (Fig. 4c, d and Supplementary Fig. 6a, Table S4).

In addition to unique pathways observed in specific interventions, a discernible fraction of pathways is enriched by both obesity and exercise interventions, indicating they are dysregulated in the obesity development and possibly reversed by exercise training. It is noteworthy that 51% of these co-enriched pathways are exclusively present in one tissue and absent in all other tissues (Supplementary Fig. 6b). MAT, pancreas and small intestine are top 3 tissues having the largest number of tissue-specific pathways co-enriched by obesity and exercise interventions. These pathways are represented by neurotransmitter receptors and postsynaptic signal transmission in MAT, VEGF signaling in pancreas, and glucose transport in small intestine, suggesting these interventions are co-regulated in specific tissues in response to both obesity development and exercise training (Supplementary Fig. 6c).

Taken together, these CMP-derived pathway analyses corroborate the above findings and point to new insights: (1) obesity remodels the multi-tissue CMP proteome landscape in a fashion substantially different from T2D whereas T2D is not an organism-wide exacerbation of the proteome response to obesity; (2) exercise training markedly yet not completely reverses the CMP proteome dysregulation by obesity, and its reversing effect varies among tissues to a large extent.

### GPCRs and RTKs dysregulated in obesity development and reversed by exercise training

Considering the essential roles of both GPCR signaling and RTK signaling in mediating energy homeostasis, we narrowed our attention to these two membrane receptor families for deepening mechanistic understanding and selecting potential therapeutic targets. From different tissues, our study identified a panel of differentially expressed GPCRs, RTKs, and protein hormones/prohormones associated with these receptors regulated in obesity, exercise, or T2D interventions (Fig. 5a). Furthermore, 5.4-60% of GPCRs and RTKs regulated in different tissues in the obesity intervention (obesity for short hereafter) showed a reverse trend of regulation responding to the exercise intervention (exercise for short hereafter), indicating the molecular adaption of these membrane receptors to exercise (Fig. 5b). We then compiled 94 GPCRs, 28 RTKs and 34 protein hormones/prohormones that are reverse-trend proteins in at least one tissue, and illustrated their tissue-specific regulation in obesity or exercise in a heat map (Fig. 5c, d).

**Figure 5.**
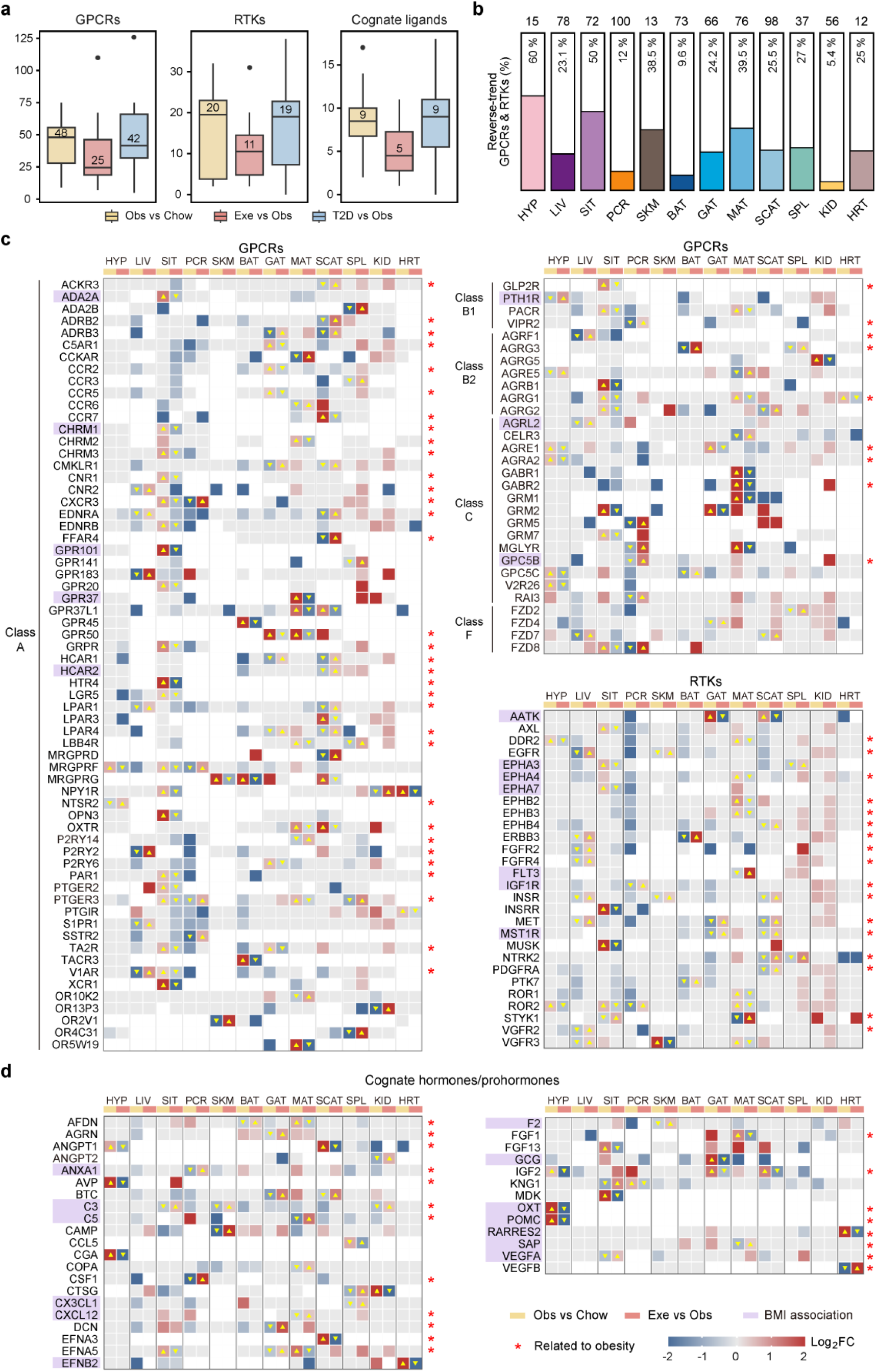
Reverse-trend GPCRs and RTKs in tissues regulated by obesity development and exercise training. (a) Number of differentially expressed GPCRs, RTKs and cognate protein ligands/prohormones identified in each tissue in the three pairwise comparisons. (b) Percentage of reverse-trend GPCRs and RTKs over all obesity-regulated GPCRs and RTKs in different tissues. Total numbers of obesity-regulated GPCRs and RTKs are annotated above the bar. (c) Heatmap showing differentially expressed GPCRs and RTKs in different tissues in the comparisons of Obs vs Chow and Exe vs Obs. Yellow triangles and color coding denote reverse-trend proteins and their trends of regulation, as defined in 4A. Red stars indicate receptors functionally linked to obesity, and purple shading marks those overlapping with BMI-associated human plasma proteins^18^ or genes^36, 37^. (d) Heatmap of differentially expressed cognate protein ligands/prohormones related to GPCRs or RTKs in different tissues, with the same denotations as in Figure 5C.

Through an extensive literature search, we found 59 reverse-trend GPCRs or RTKs to be previously disclosed as regulators of obesity-related metabolic dysfunction or energy imbalance and serve as approved or promising anti-obesity targets. For example, in the GPCR families, β-2 adrenergic receptor (β2AR) in SCAT and β-3 adrenergic receptor (β3AR) in SCAT and GAT were all down-regulated in obesity and up-regulated in exercise. This result is in line with the roles of both β2AR and β3AR in simulating lipolysis in WAT and the additional role of β3AR in promoting “browning” of WAT to enhance energy expenditure^31, 64, 65^. Thus, selective agonists of either receptor are actively explored for the treatment of obesity and related metabolic syndrome^31^. Cannabinoid receptor 1 (CB1) in small intestine was up-regulated in obesity and reversed in exercise whereas cannabinoid receptor 2 (CB2) in liver was down-regulated in obesity and reversed in exercise. Consistently, overactivation of peripheral CB1 is known to influence hormone secretion, promote lipid storage, and disrupt metabolic homeostasis, implicating CB1 antagonism as a therapeutic strategy for body weight control^66, 67^. Also supporting our data, CB2 activation mitigates inflammation in peripheral tissues and improve insulin sensitivity, making CB2 selective agonists promising for treating obesity-related metabolic disorders^31, 68^. Neurotensin receptor type 2 (NTSR2), a neuropeptide receptor relevant to appetite control, was down-regulated in hypothalamus in obesity and reversed in exercise, supporting that impaired neurotensin signaling in hypothalamus contributes to hyperphagia and weight gain^69, 70^.

In the RTK family, a panel of growth factor receptors including EGFR, FGFR2/4, VEGFR2/3, MET and the aforementioned insulin receptor were all down-regulated in liver in obesity and reversed in exercise. Concordantly, FGFR2/4 in liver play major roles in hepatic lipid metabolism, insulin sensitivity, or bile acid regulation, and their disrupted activities contribute to hepatic steatosis, inflammation, and insulin resistance^71, 72^. EGFR, VEGFR, and MET also play significant roles in hepatic glucose and lipid metabolism, with their dysregulation contributing to obesity-associated liver pathologies and insulin resistance^73, 74, 75^. Given that activation of these receptors correlates with insulin resistance and hepatic lipid accumulation, their down-regulation in obesity may implicate compensatory mechanisms to counteract metabolic disruption. Therefore, pharmacological modulators of these growth factor receptors are being evaluated in clinical trials to treat obesity-associated liver diseases by restoring glucose and lipid metabolism^30, 76^.

On the other side, 22 GPCR/RTK-associated protein hormones or prohormones with a reverse trend of regulation are also implicated in energy homeostasis and obesity development. Particularly, a few growth factors including FGF1, VEGFB, and insulin-like growth factor 2 (IGF2) were reversely regulated in heart or adipose tissues responding to obesity and exercise, in line with their major roles in mediating insulin sensitivity, lipid oxidation and fat accumulation^77, 78, 79^. In addition, prohormones for several appetite-regulating neuropeptides were up-regulated in hypothalamus in obesity and reversed in exercise. Among them, elevated vasopressin (AVP) is known to amplify chronic stress-induced hyperphagia while oxytocin (OXT) and pro-opiomelanocortin (POMC) both can promote satiety and reduce food intake^80, 81, 82^. Thus, obesity-induced increase of OXT and POMC may imply a resistance mechanism resulting from weakened anorexigenic signals and energy imbalance in hypothalamus. Altogether, by systematically mining these reverse-trend membrane receptors and their cognate ligands, we demonstrate the power of our CMP proteome profiling strategy to uncover the organism-wide and tissue-specific co-regulatory network that is extensively remodeled in obesity development or exercise training. Alignment of the dysregulation of many reverse-trend proteins with their functional implications in obesity not only pinpointed the disease-preventing benefits of exercise but also prompted us to investigate the potential of unrelated CMPs as new anti-obesity targets.

### Discovery of hypothalamic PTHRs as novel GPCR regulators of feeding, body weight, and exercise-induced adaptation

The hypothalamus is a critical integrative center for the regulation of energy homeostasis^83, 84, 85^. To uncover previously unrecognized regulators of energy balance within the hypothalamus, we focused on PTH1R, a GPCR whose protein abundance was significantly down-regulated in obese mice compared to chow-fed controls and restored to baseline following an exercise intervention (Fig. 6a, Table S3-HYP). PTH1R belongs to the parathyroid hormone (PTH) receptor subfamily, which also includes PTH2R. Both human PTH1R and PTH2R respond to the endogenous ligand PTH, and human PTH2R is selectively activated by the tuberoinfundibular peptide of 39 residues (TIP-39), also known as PTH2^86^. For PTH2R which was originally undetected in our CMP proteome profiling, we developed a more specific and sensitive PRM-MS assay to show its protein abundance was slightly reduced in obese mice and significantly elevated after exercise training (Supplementary Fig.7). Additionally, its ligand TIP-39 was down-regulated in obese mice and kept unchanged after exercise training (Fig. 6b, Table S3-HYP).

**Figure 6.**
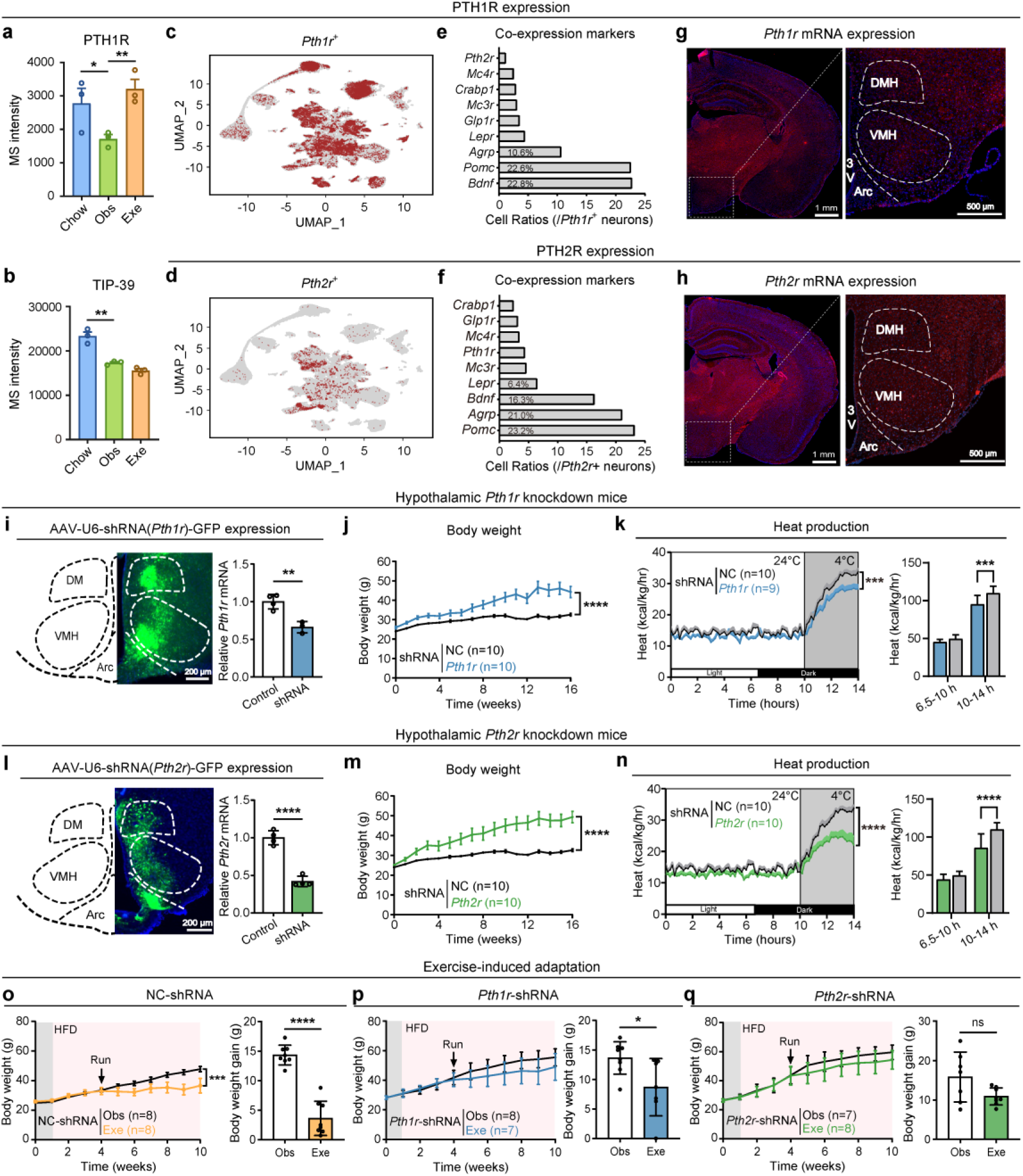
Hypothalamic parathyroid hormone receptors (PTHRs) regulate feeding, body weight, and exercise-induced adaptation. (a-b) MS-based relative quantification of PTH1R (a) and TIP-39 (b) in mice from Chow, Obesity (Obs), and Exercise (Exe) groups in biological triplicate. (c-d) Uniform manifold approximation and projection (UMAP) visualization of *Pth1r*⁺ cells (c) and *Pth2r*⁺ cells (d) in HypoMAP. Each point represents a single cell. (e-f) The ratios of *Pth1r*⁺ (e) and *Pth2r*⁺ (f) neurons co-expressing *Mc4r, Crabp1, Mc3r, Glp1r, Lepr, Agrp, Pomc*, or *Bdnf*. (g-h) Hypothalamic expression of *Pth1r* (g) and *Pth2r* (h) mRNA in *C57* mice. (i) Representative expression of AAV-U6-shRNA(*Pth1r*)-GFP (left) and reduction of *Pth1r* mRNA (right) expressed in the hypothalamus of *Pth1r*-shRNA mice. Scale bars, 200 mm. The scrambled shRNA group was used as the control. (j-k) Changes in body weight (j) and heat production (k) of *Pth1r*-shRNA mice. (l) Representative expression of AAV-U6-shRNA(*Pth2r*)-GFP (left) and reduction of *Pth2r* mRNA (right) expressed in the hypothalamus of *Pth2r*-shRNA mice. Scale bars, 200 mm. The scrambled shRNA group was used as the control. (m-n) Changes in body weight (m) and heat production (n) of *Pth2r*-shRNA mice. (o-q) Changes in body weight induced by exercise intervention in control mice (o), *Pth1r*-shRNA mice (p) and *Pth2r*-shRNA mice (q). The column graph displays the comparative weight gain changes after exercise intervention. Data are presented as mean ± SEM and analyzed by two-way RM ANOVA or t test. \**p* < 0.05, \*\**p* < 0.01, \*\*\**p* < 0.001 *vs* corresponding control group. ns, not significant.

**Figure 7.**
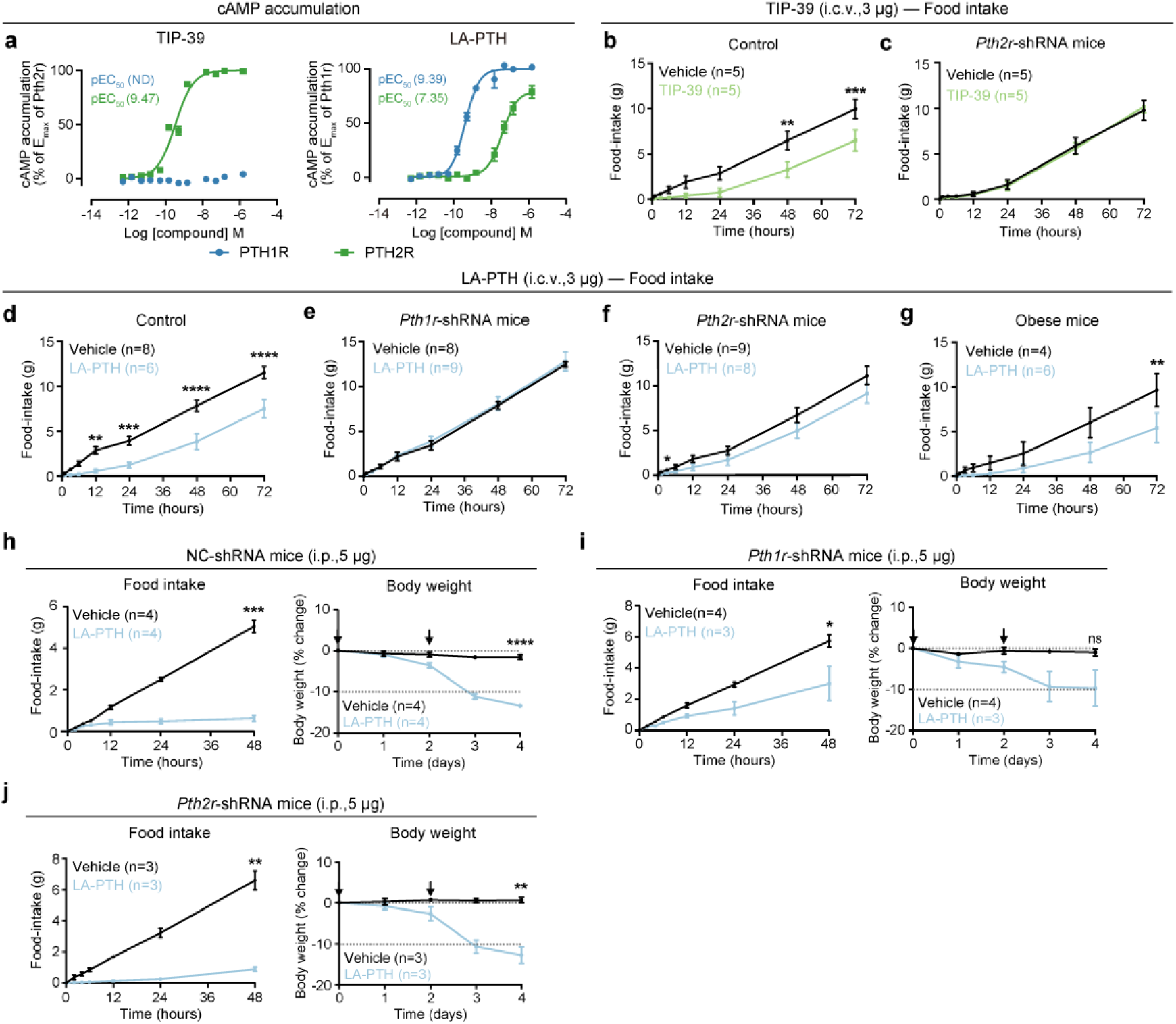
Agonists targeting hypothalamic PTHRs mediate feeding and body weight. (a) Dose-response curves of cAMP accumulation in HEK293T cells expressing mouse PTH1R or PTH2R treated with agonist LA-PTH or TIP-39. Mean pEC_50_ values from three independent experiments are shown (see Supplementary Fig. 8c, d for complete data). ND: not determined. (b-c) Changes in food intake of *C57* mice (b) and *Pth2r*-shRNA mice (c) after intracerebroventricular (i.c.v.) TIP-39 (3 μg) at the indicated time. (d-g) Changes in food intake of *C57* mice (d), *Pth1r*-shRNA mice (e), *Pth2r*-shRNA mice (f) and obese mice (g) after central administration of LA-PTH (3 μg). (h-j) Cumulative food intake and body weight changes in control mice (h), *Pth1r*-shRNA mice (i) and *Pth2r*-shRNA mice (j) following intermittent intraperitoneal (i.p.) injections of LA-PTH (5 μg, every other day) under a HFD condition. Data are presented as mean ± SEM and analyzed by two-way RM ANOVA or t test. \**p* < 0.05, \*\**p* < 0.01, \*\*\**p* < 0.001 *vs* corresponding control group. ns, not significant.

To define the molecular identity of PTHR-expressing cells, we integrated multiple single-cell RNA-sequencing (scRNA-seq) data sets encompassing 384,925 mouse hypothalamic cells reported previously^84^. We identified 17,204 *Pth1r* and 2,717 *Pth2r*-expressing cells, with the vast majority being neuronal (9,486 and 2,652 cells, respectively) (Fig. 6c, d). Co-expression analysis revealed that a substantial proportion of *Pth1r*+ neurons expressed *Bdnf* (22.8%), *Pomc* (22.6%), or *Agrp* (10.6%) (Fig. 6e). Similarly, *Pth2r*+ neurons frequently co-expressed *Pomc* (23.2%), *Agrp* (21.0%), *Bdnf* (16.3%), or *Lepr* (6.4%) (Fig. 6f). Conversely, canonical energy-balance receptors (e.g., *Mc4r*, *Mc3r*, *Glp1r*) were detected in fewer than 5% of *Pthr*+ cells. This low co-expression rate was also observed with the exercise-induced gene *Crabp1*^87^, and the population of neurons co-expressing both *Pth1r* and *Pth2r* was minimal (Fig. 6e, f). These findings suggest that hypothalamic PTHR signaling defines neuronal populations largely distinct from established metabolic pathways. We then performed spatial mapping *via* RNAscope *in situ* hybridization to reveal the localization of *Pth1r* and *Pth2r* transcripts to distinct hypothalamic nuclei: *Pth1r* was sparsely distributed across the hypothalamus, whereas *Pth2r* showed prominent expression in the mediobasal hypothalamus (MBH), including ventromedial hypothalamus (VMH) and arcuate nucleus (ARC) (Fig. 6g, h).

To assess the physiological roles of hypothalamic PTHRs, we selectively knocked down each receptor in WT mice *via* stereotactic delivery of AAV-encoded shRNAs targeting *Pth1r* or *Pth2r* (hereafter referred to as *Pth1r*-shRNA mice or *Pth2r*-shRNA mice). This intervention reduced hypothalamic *Pth1r* and *Pth2r* mRNA levels by 34% and 58%, respectively, compared to control mice (Fig. 6i, l). Strikingly, *Pth1r*-shRNA mice exhibited progressive weight gain, increased adiposity, decreased lean mass, and impaired cold-induced thermogenesis (Fig. 6j, k and Supplementary Fig. 8a), underscoring a critical role of hypothalamic PTH1R in energy balance. Furthermore, hypothalamic knockdown of *Pth2r* recapitulated this metabolic phenotype, including progressive obesity, increased adiposity, lean mass loss, and blunted thermogenic responses (Fig. 6m, n and Supplementary Fig. 8b). These results indicate that both PTH1R and PTH2R are non-redundant regulators of energy balance within the hypothalamus.

Given that the PTHR system exhibited a reverse-trend response to obesity progression *vs* exercise training, we set out to determine whether PTHR signaling contributes to the exercise-induced adaptation to metabolic stress. While feeding *Pth1r*-shRNA and *Pth2r*-shRNA mice with HFD, we subjected them to a standardized exercise regimen in the same manner as specified in Figure 3a. In the control group, endurance exercise reduced the body weight gain of HFD-fed mice to 25.3% of that in the sedentary counterpart (Fig. 6o). Remarkably, both *Pth1r*-shRNA and *Pth2r*-shRNA mice exhibited a markedly attenuated weight loss, with post-exercise body weight gains remaining at 63.5% and 68.9% of the sedentary levels respectively (Fig. 6p, q). These findings indicate that hypothalamic PTHR signaling via two receptors plays a critical role in exerting the benefits of exercise in suppressing weight gain and metabolic dysfunction under HFD-induced metabolic stress.

### Targeting hypothalamic PTHR signaling to combat obesity

To evaluate the therapeutic potential of hypothalamic PTHR activation in obesity, we first assessed whether central administration of selective ligands could modulate the feeding behavior. TIP-39-a high-affinity, sub-nanomolar agonist for mouse PTH2R-robustly stimulated Gs-mediated cAMP accumulation *in vitro*, consistent with its known activity at human PTH2R (Fig. 7a and Supplementary Fig. 8c, d).^88^ Intracerebroventricular (i.c.v., third ventricle) delivery of TIP-39 significantly suppressed acute food intake in lean *C57BL/6J* mice, an effect that was completely abolished in *Pth2r*-shRNA mice, suggesting PTH2R was the primary target of TIP-39 in apetitie control (Fig. 7b, c).

We also tested LA-PTH, a long-acting PTH analog^89^ that potently activates mouse PTH1R and exhibits weaker activity at PTH2R (Fig. 7a). Central administration of LA-PTH similarly reduced food intake in *C57BL/6J* mice (Fig. 7d). Notably, the anorexigenic effect of LA-PTH were partially attenuated in *Pth2r*-shRNA mice and entirely abrogated in *Pth1r*-shRNA mice (Fig. 7e, f). Importantly, this anorexigenic response was preserved in HFD-induced obese (DIO) mice, and LA-PTH elicited a more pronounced effect than TIP-39 (Fig. 7g and Supplementary Fig. 7e), suggesting that PTH1R activation may be particularly effective in the context of established obesity. To translate these findings toward therapeutic application, we evaluated whether peripheral delivery of LA-PTH could recapitulate the anti-obesity effects observed with central administration. Indeed, intraperitoneal (i.p.) injection of LA-PTH led to a dose-dependent reduction of the body weight of DIO mice, with 5 μg being an effective minimal dose compared to 3 μg, 10 ug or 30 ug injections (Supplementary Fig.7f). Peripheral injection with 5 μg LA-PTH did not affect the locomotor activity (Supplementary Fig.7g). Critically, the metabolic effects of LA-PTH were abrogated in DIO mice with hypothalamic *Pth1r* knockdown (Fig. 7h, i). In contrast, the response was only partially attenuated in *Pth2r*-shRNA mice (Fig. 7j), demonstrating a selective requirement for PTH1R in mediating the systemic benefits of LA-PTH. Although earlier studies reported a similar weight loss phenotype by systemic administration of PTH or PTH-related hormones to DIO mice, these effects were all attributed to PTH1R expressed in liver and adipose tissues^90, 91^. The contribution of the hypothalamic PTHR system to weight control and energy metabolism has not been recognized. Collectively, our data establish hypothalamic PTHRs-particularly PTH1R-as a druggable node for the central control of energy balance and highlight the potential of peripherally administered PTH analogs as a promising strategy for obesity treatment.

## Discussion

Complementary to all previously reported global proteome maps of human or animal tissues^14, 16, 17, 92^, our study establishes a whole-organism molecular atlas focusing on cell membrane proteins which encompass the largest number of known druggable targets compared to proteins in other cellular compartments. Through adapting our CMP proteome profiling approach to multiple tissues, we built the most comprehensive CMP proteome map across 22 mouse tissues with largely improved identification depth and quantification consistency for CMPs (Fig. 1). Of note, compared to a seminal work from the Kuster group presenting a global proteome map across 41 mouse tissues which integrated MS data from 1,312 acquisitions^16^, our study which combined MS data from 85 acquisitions enabled additional mapping of 584 new CMPs including 100 new GPCRs, illustrating the advantage of our strategy in profiling low-abundance membrane proteins on the cell surface. In addition, our study describes the first multi-tissue map of endogenous protein ligands or prohormones associated with members in the GPCR superfamily (Fig. 2d), which could be used to infer the potential activation of certain GPCRs upon binding to their cognate ligands. However, whether these receptors paired with their ligands undergo activation and trigger downstream signaling also depends on other factors (*e.g.* the concentration and conformation of their ligands in specific tissues), which warrants further investigation.

By mapping the multi-tissue proteome dynamics in mouse models of obesity, T2D and exercise, our study systematically characterized the CMP proteome remodeling in response to metabolic disease progression *versus* endurance exercise training. Altogether, we identified 2,303 and 2,216 differentially expressed CMPs from 12 examined tissues responding to obesity development and exercise training, respectively-approximately 20 times more than previously reported in comparable omics studies^8, 9^. Specifically, a previous multi-omics study of 27 tissues from monkeys reported 98 CMPs significantly regulated in the development of obesity^8^, and another comprehensive study of 19 tissues from rats documented 80 CMPs regulated by 8-week exercise training^9^, based on data re-analysis with the same criteria for DEP selection (see Methods). It is noteworthy that the substantial discrepancy in the identification of DEPs from the proteome data *vs* DEGs from the transcriptome data as observed in our work was also mentioned in the two previous studies^8, 9^.

With the unprecedented depth and accuracy of CMP profiling, this study can be leveraged to advance our understanding of CMP-involved mechanisms underpinning obesity/T2D pathogenesis *versus* exercise adaptation. Interestingly, within individual tissues, pairwise comparisons of obesity vs chow, exercise vs obesity, and T2D vs obesity yielded a large proportion of DEPs (39.1%-79.2%) and DEP-enriched pathways (38.5%-91.7%) that are unique to a single comparison (Fig. 4c and Supplementary Fig. 5a, 6a). This indicates that the development of obesity and its progression to T2D had distinct effects on CMP proteome homeostasis, which were distinguished from exercise-induced effects. More intriguingly, we observed a number of GPCRs, RTKs and glucose transporters with critical roles in mediating metabolism and energy balance were dysregulated in obesity development and reversed by exercise training in different tissues (Fig. 4a, 5c). This result illustrates the organism-wide and tissue-specific rewiring of the key molecular network by exercise training, which could lead to the alleviation or prevention of obesity and related metabolic disorders.

Our CMP-focused molecular profiling in metabolic disease and exercise models led to the identification of parathyroid hormone receptors PTH1R and PTH2R in the hypothalamus as novel regulators of feeding and energy metabolism in the central nervous system (CNS). Notably, PTH1R primarily expressed in bone and kidney regulates calcium homoeostasis and skeleton development, and it serves as a clinically proven target for hypoparathyroidism and osteoporosis^93^. PTH1R in the CNS has been implicated in protection against neuroinflammation and neurodegeneration, with positive effects on memory and hyperalgesia^94, 95^. Moreover, PTH2R predominantly expressed in the brain has been reported to modulate auditory and nociceptive processes, and sexual maturation^94, 96^. However, the metabolic functions of neither receptors in the CNS remain largely unexplored. Clinically, weight gain following parathyroidectomy suggests that the PTH-PTHR axis plays an important role in body weight regulation^97^. Previous studies attributed the body weight - lowering effects of PTH/PTHrP primarily to the promotion of adipocyte browning^91, 98^. Here, we demonstrate that targeting hypothalamic PTHR signaling potently ameliorates obesity in mice. Moreover, peripheral administration of the long-acting PTH analogue (LA-PTH) robustly suppresses appetite and reduces body weight, an effect mediated by hypothalamic PTHRs. Altogether, through pharmacological and genetic interventions, we uncovered the unknown roles of the hypothalamic PTH1R/PTH2R system in regulating food intake and energy balance. Notably, this pathway also contributes to exercise-induced weight loss, revealing a novel mechanistic link between central PTHR signaling and physiological stress responses. Thus, we demonstrated the capability of our proteomic strategy in the identification of tissue-specific key regulators, which could inform therapeutic interventions to tackle metabolic disorders. In addition, the remaining reverse-trend GPCRs or RTKs with no implications in obesity or T2D would deserve future investigations.

Limitations of the current study include not considering the sex differences in developing the models of metabolic diseases and exercise training, and using mouse models to mimic the disease progression and exercise adaptation. The translational potential of this study, though limited by the mouse models, is demonstrated by the concordance of a portion of our CMP regulation data with human plasma proteome and GWAS data as well as the recapitulation of a plethora of known regulators of obesity-related phenotypes. In the future, our CMP profiling approach can be employed to study the multi-tissue proteomic responses of male and female nonhuman primates to obesity development *versus* exercise training. Moreover, our current approach to profile the CMP proteome in heterogeneous tissues lacks cell type specificity and spatial localization. We envision that this drawback could be in part overcome by integrating our proteomics data with single-cell and spatial transcriptomics data to derive the distribution of CMPs in specific cell types or clusters (Fig. 6c-f). Finally, a more comprehensive *in vivo* characterization of central PTHR signaling is required to elucidate its mechanism in regulating energy balance particularly in the hypothalamus.

In summary, our study presents an enabling platform to map the tissue distribution and dynamics of the CMP proteome together with the associated protein hormones and prohormones. Application of this approach to the study of proteomic responses to metabolic disease development *vs* exercise training provided new biological insights and led to the discovery of novel GPCR regulators of diet-induced obesity and exercise-mediated adaptation. We anticipate this unique CMP omics data resource would provide new opportunities to understand the interplay between metabolic disease progression and exercise-induced benefits that are both regulated by cell membrane proteins.

## EXPERIMENTAL DESIGN AND SAMPLE PREPARATION

### Mice and mouse models

Adult male C57BL/6J (wt) mice (10-week-old) were purchased from GemPharmatech Co., Ltd. (Nanjing, China). For the mouse model development, mice were randomly assigned into four groups: chow, obesity (Obs), exercise (Exe), and type 2 diabetes (T2D). The Chow group was fed with standard laboratory chow food during the entire experimental period. The Obs group was fed with high-fat diet (HFD; D12492, Research diets, USA) for 9 consecutive weeks to cause diet-induced obesity. The Exe group was fed with HFD and underwent treadmill exercise training (training procedure described below). The T2D group was fed with HFD for 13 consecutive weeks to cause T2D related phenotypes.

Mice were housed under constant temperature and humidity in a 12h light/12h dark cycle (light, 9 PM to 9 AM) and provided with ad libitum (ad-lib) access to water. All animal procedures were performed in strict accordance with the Animal Research: Reporting of In Vivo Experiments Guidelines. The animal experiments were approved by the Laboratory Animal Management Committee of ShanghaiTech University.

### Treadmill exercise training

Mice underwent a treadmill exercise using a motorized system (TSE Systems, Berlin, Germany). After 2 weeks of HFD feeding, mice underwent a one-week acclimation phase involving daily treadmill running (30 min/day at 10 m/min). From weeks 4–9 of HFD feeding (6 weeks total), exercise training was conducted: 1-hour treadmill sessions, five days per week, at 12 m/min with a 21-second acceleration ramp. Mice in the exercise (Exe) group were euthanized 24 hours after the final session.

### Tissue collection

At the study endpoint, the tissues including adrenal gland (AD), brain (BRN), hypothalamus (HYP), heart (HRT), kidney (KID), liver (LIV), lung (LNG), brown adipose tissue (BAT), mesenteric adipose tissue (MAT), gonadal adipose tissue (GAT), subcutaneous adipose tissue (SCAT), skeletal muscle (SKM), pancreas (PCR), prostate (PRS), skin (SKN), small intestine (SIT), spleen (SPL), stomach (STM), testis (TES), thymus (THM), eye (EYE), tongue (TON), and urinary bladder (UB) were dissected for CMP proteomics and RNA-seq experiments. Twenty-two (all the above except HYP) were dissected from normal mice and twelve tissues (HYP, LIV, SIT, PCR, SKM, BAT, GAT, MAT, SCAT, SPL, KID, HRT) from each group of mouse models.

For normal mice, independent experimental replicates for most tissues were collected from individual mice. Exceptions were AD, UB, and EYE, for which each replicate was prepared by pooling tissues from three mice. All tissues were collected in four independent replicates except for AD, UB, and EYE in three replicates. For each experimental group of the model models, independent replicates for most tissues were also collected from individual mice, with three replicates per tissue. The hypothalamus was an exception as each replicate was prepared by pooling tissues from three mice.

### Intraperitoneal glucose tolerance tests (ipGTTs)

Intraperitoneal glucose tolerance tests (ipGTTs) were conducted after 12h overnight fasting following an intraperitoneal (i.p.) injection of glucose (20% dextrose) at a dose of 1.0 g/kg body weight. Glucose measurement was determined by a hand-held glucometer (Accu-Chek Performa Connect, Roche, Switzerland). Blood samples were collected from the tail vein at the indicated time.

### Cell membrane protein extraction

Tissues were homogenized in isolation buffer containing 30 mM Tris-HCl (pH 7.6), 0.1 mM EDTA, 300 mM sucrose, 0.5% bovine serum albumin (BSA), and EDTA-free complete protease inhibitor cocktail tablets (Thermo Scientific). The homogenate was centrifuged at 2,500 × g for 10 min at 4°C, and the supernatant was further centrifuged at 10,000 × g for 20 min at 4°C. The supernatant was collected for ultracentrifugation at 160,000 x g for 1 hour at 4°C. The resulting membrane pellet was sequentially washed with 1 M KCl, 100 mM Na_2_CO_3_, and 100 mM Tris-HCl (pH 7.6), followed by ultracentrifugation at 160,000 × g for 1 hour at 4°C. The final pellet was resuspended in a lysis buffer containing 5% SDC and 50 mM NH_4_HCO_3_, and heated at 95°C for 5 min. Protein concentration was determined using a BCA assay (TIANGEN, China).

### Cell membrane protein digestion and high-pH reversed-phase fractionation

About 10 μg protein extract from each tissue replicate for normal mice or mouse model groups were reduced with 15 mM dithiothreitol (DTT) at 56°C for 40 min, alkylated with 40 mM iodoacetamide (IAA) at room temperature for 30 min, and further treated with 25 mM DTT at 37°C for 30 min. After diluting SDC to 0.5% with 50 mM NH_4_HCO_3_, proteins were digested with rLys-C (Beijing Shengxia Proteins Scientific Ltd., China) at an enzyme-to-protein ratio of 1:100 (w/w) for 3 hours at 37°C, followed by overnight digestion with trypsin (Promega) at an enzyme-to-protein ratio of 1:50 (w/w) at 37°C. Digestion was quenched with 1% trifluoroacetic acid (TFA), and the protein digest was desalted using C18 microspin cartridges (Omicsolution, China), lyophilized under vacuum and stored at -80°C.

For protein digests from tissues of mouse models, half digests from all replicates of each tissue from four experiment groups were pooled and loaded onto an equilibrated, high-pH reverse-phase fractionation spin column (Pierce). The protein digests were bound to the hydrophobic resin, and desalted via water washing. A stepwise gradient of acetonitrile (4-50%) in a volatile high-pH elution buffer was then applied to fractionate the peptides into eight sequential eluates. All the fractions were dried by vacuum centrifugation and stored at -80°C until nano-liquid chromatography-tandem MS (nanoLC-MS/MS) analysis.

### NanoLC-MS/MS data acquisition

For normal mice, the protein digest from each replicate of each tissue (22 tissues altogether) was subjected to nanoLC-MS/MS analysis in the DIA mode. Peptide samples were separated on an analytical column (300 mm x 75 μm) and in-house packed with C18-AQ 1.9-μm C18 resin (Dr. Maisch GmbH), by a 130-min gradient of 4-8% solvent B (0.1% formic acid in 80% acetonitrile) for 20 min, 8-22% solvent B for 70 min, 22-45% B for 30 min, 45-100% B for 4 min, 100% B for 4 min, 100-4% B for 30s and 4% B for 90s at a flow rate of 300 nl/min. Peptides were analyzed using an Easy-nLC 1200 system coupled to a Orbitrap Eclipse Tribrid mass spectrometer equipped with a FAIMS Pro interface (Thermo Fisher Scientific). The instrument was operated in the positive-ion NSI mode. Typically, electrospray ionization was carried out with a spray voltage of 2200 V. Two FAIMS compensation voltages (CV) of -45 V and -60 V allowed the comprehensive acquisition of all peptides. In the DIA mode, the 42 variable windows were set to cover a range of 350 to 1500 m/z with the following settings: resolution of Orbitrap analyzer, 60,000 for MS1 and 30,000 for MS2; AGC, 3e6 in MS1 and 1e5 in MS2; maximum injection time, 50 ms in MS1 and 40 ms in MS2. The stepped collision energy was set to 32 ± 5%.

For each experimental group of the mouse models, the protein digest from each replicate of each tissue (12 tissues altogether) was split into two, one half for DIA MS and the other half for peptide pre-fractionation and DDA MS analysis. Peptide samples were analyzed using a nanoElute LC system coupled to a timsTOF Pro 2 mass spectrometer (Bruker). Peptides were separated on the same analytical column using a 90 min-gradient of 3-22% solvent B (0.1% formic acid in acetonitrile) for 65 min, 22-37% B for 15 min, 37-80% B for 5 min and 80% B for 5 min at a flow rate of 300 nl/min. DIA MS data were acquired in the DIA parallel accumulation-serial fragmentation (PASEF) mode. The capillary voltage was set to 1,500 V, mass range for MS scans was 100 to 1,700 m/z, ion mobility scanning was from 0.75 to 1.4 Vs cm^−2^, and the ramp time was set to 100 ms. Isolation windows of a width of 25 m/z were defined across the mass range of 452 to 1,152 m/z. The collision energy was ramped linearly based on ion mobility from 59 eV at 1/K0 = 1.6 Vs cm^−2^ to 20 eV at 1/K0 = 0.6 Vs cm^−2^. DDA MS data of all the fractions were acquired on timsTOF Pro 2 mass spectrometer (Bruker) in the DDA PASEF mode with 10 PASEF MS/MS scans. The capillary voltage was set to 1,500 V and the mass range for MS scans was set to 100-1,700 m/z with ion mobility scanning from 0.6 to 1.6 V s cm^−2^. The ramp time was set to 100 ms and total cycle time was 1.17 s. Precursors with 0-5 charge state were selected with target intensity of 20,000 and intensity threshold of 2,500 for fragmentation. The dynamic exclusion was set to 0.4 min and the isolation width was set to 2 m/z for m/z < 700 and 3 m/z for m/z > 800. The collision energy was also ramped linearly as a function of the mobility from 59 eV at 1/K0 = 1.6 Vs cm^−2^ to 20 eV at 1/K0 = 0.6 Vs cm^−2^.

### Parallel Reaction Monitoring (PRM)-MS assay

For CMP digests separately prepared from hypothalamic tissues collected from the chow, obesity, and exercise groups, peptides were separated on the same analytical column with the same gradient as the DIA MS analysis. PRM data were acquired on a timsTOF Pro 2 (Bruker) in the PRM-PASEF mode, using a scheduled target list of proteotypic peptides of mouse PTH2R that were predicted using DIA-NN. Auto-calibration was performed before each run, with high sensitivity detection mode enabled. QC injections of a HeLa digest confirmed the mass accuracy (< 10 ppm). Each group was prepared in experimental triplicate with randomized injections.

Raw files were converted using MSConvert (ProteoWizard), and processed in Skyline (v25, MacCoss Lab)^99^ mostly with default settings. Library ion matching tolerance was set to 10 ppm, and retention time filtering was applied to include all matching scans. The cutoff of isotope dot product (idotp) indicating experimental and library spectral matching was 0.85. The integrated peptide intensity was used for differential analysis.

### Tissue RNA extraction and RNA-sequencing

For RNA-sequencing analysis, two biological replicates were prepared from each tissue of the normal mice, and three biological replicates were collected from each tissue of each experimental group for the mouse models. Each replicate was from an individual mouse. Total RNA was extracted from each tissue sample and treated with deoxyribonuclease I (DNase I). Messenger RNA was purified from total RNA using poly-T oligo-attached magnetic beads. After fragmentation, the first strand cDNA was synthesized using random hexamer primers followed by the second strand cDNA synthesis. The library was ready after end repair, A-tailing, adapter ligation, size selection, amplification, and purification. Quality check of the library was performed with Qubit and real-time PCR for quantification and with bioanalyzer for size distribution analysis. Library preparation and RNA sequencing were outsourced to Genewiz (Suzhou, China) and Novogene (Beijing, China).

### Intracellular cAMP accumulation assay

Intracellular cAMP was quantified using the cAMP-Gs Dynamic Kit (Cisbio) in a homogeneous time-resolved fluorescence (HTRF) format. HEK293T cells were seeded into 384-well shallow plates at 3,000 cells per well in DMEM containing 1% dialyzed FBS. For stimulation, cells transfected with mouse Pth1r or Pth2r were incubated for 20 min at room temperature with serial dilutions of LA-PTH (Changzhou Kanglong Biotech, China) or human TIP-39 (Nanjing Jietai Biotech, China) prepared in assay buffer (1× HBSS + 0.1% BSA). Following ligand stimulation, anti-cAMP-cryptate and cAMP-d2 conjugate were added simultaneously, and reactions were allowed to proceed for 60 min at room temperature. HTRF signals were measured at 665 nm and 620 nm on an EnVision plate reader (PerkinElmer). Dose–response curves were fitted to a four-parameter logistic model in GraphPad Prism 8.2.1, and responses were normalized to the maximal LA-PTH induced cAMP accumulation.

### Flow Cytometry

Surface expression of 3×Flag-tagged mouse PTH1R and PTH2R was determined by flow cytometry. HEK293T cells expressing the receptors were harvested by centrifugation, washed, and incubated with anti-FLAG M2-FITC antibody (Sigma, F4049; 1:1,000) for 20 min at 4 °C. Fluorescence was recorded on a CytoFLEX flow cytometer (Beckman Coulter) and data were analyzed using FlowJo v10 (BD Biosciences).

### RNA in situ hybridization

For Pth1r/Pth2r RNA in situ hybridization, mice were anesthetized and perfused with 4°C pre-chilled DEPC-PBS. Brains were removed, fixed in 4% PFA for 16 hours, and dehydrated in 15% sucrose solution for one day followed by 30% sucrose solution for three days. Brain tissue sections (16 μm thick) were prepared using a cryostat microtome. Hybridization was performed using the RNAscope 2.5HD Kit (#322350, Advanced Cell Diagnostics) according to the manufacturer’s instructions. Probes for Pth1r (#426191, Advanced Cell Diagnostics) and Pth2r (#543351, Advanced Cell Diagnostics) were designed by Advanced Cell Diagnostics, with the DapB probe serving as a negative control.

### AAV vectors

AAV vectors (with a titer of >10^12^) carrying specific genes were packaged by Shanghai Sunbio Biotechnology Co. (Shanghai, China), including AAV2/8-U6-shRNA (*Pth1r*)-EF1a-eGFP-pA and AAV2/8-U6-shRNA (*Pth2r*)-EF1a-eGFP-pA. The target sequences for these genes are GCTACCAACTACTACTGGATT (*Pth1r*), CTGGTATTTGGCGTGCATTAC (*Pth2r*).

### Stereotaxic injection

Stereotaxic surgery was performed under deep anesthesia using isoflurane, and mice were placed in a stereotaxic apparatus (David Kopf Instruments, #PF-3983; RWD Life Science, #68030; Thinker Tech Nanjing Biotech, #SH01A). Two injection sites were selected per hemisphere to ensure the targeted delivery within the hypothalamus. The stereotaxic coordinates, based on the Paxinos & Franklin mice brain coordinates (3rd edition), were as follows: ML: ±0.4 mm, AP: -1.1 mm, DV: -5.65 mm; ML: ±0.4 mm, AP: -1.1 mm, DV: -5.95 mm. At each site, 150 nl of viral solution was injected using a pulled-glass pipette and a pencil type oil hydraulic injector (MO-10, Tritech Research, USA) with customized manipulators at a constant rate of 30 nl/min. The needle was left in place for additional 5 minutes following injection to minimize backflow before slow withdrawal. A feedback heater was used to keep mice warm during surgeries. Mice were monitored post-operatively and allowed to recover for at least 7 days before subsequent experimental procedures.

### Food intake, body weight, and body composition analysis

Body weight and food intake were continuously monitored under standard housing conditions. To assess changes in body composition, quantitative magnetic resonance (QMR) spectroscopy was performed using a Bruker LF50 Body Composition Analyzer (Bruker, Karlsruhe, Germany). Fat mass and lean mass were quantified and normalized to total body weight for comparative analysis.

### Indirect calorimetry

Heat production was monitored with a comprehensive lab animal monitoring system (CLAMS, Columbus Instruments, USA). Mice were individually housed in metabolic cages with ad libitum access to food and water and allowed to acclimate for 10 hours prior to data collection. The experiment was conducted at an ambient temperature of 24°C, with animals subjected to a 4°C cold challenge for 4 hours following the acclimation period. Data were collected at 10-minute intervals.

During the study, mice were maintained on a 12-hour light/dark cycle, with lights on at 21:00 and off at 09:00. The effect of hypothalamic Pth1r and Pth2r knockdown on energy expenditure was further evaluated by comparing the area under the curve (AUC) of energy expenditure before and after cold exposure.

### Intracerebroventricular cannulation and treatment

In the food intake assay, mice were surgically implanted with the cannula (RWD Life Science, 62002, 62102) in the third ventricle (3V) (ML: 0.0 mm, AP: -1.25 mm, DV: - 5.30 mm), and the cannula was secured to the skull with dental cement (C&B Metabond Quick Adhesive Cement System). All cannulated mice were individually caged post-surgery for 7–10 days’ recovery with chow food and water.

To reduce stress-induced variability in feeding behavior, mice were gently handled daily for one week before formal administration. During this habituation period, each mouse received three intracerebroventricular (3V) injections of sterile saline (2 μl per injection) on alternating days via micro syringe through the cannula. This pre-treatment step was designed to acclimate the animals to the injection procedure and minimize the potential confounding effects of handling or injection-related stress. Following acclimation, the test compound (LA-PTH or TIP-39) was administered via 3V injection at a dose of 3 μg in 2 μl, delivered in the morning to minimize circadian variability. After each injection, the needle was left in place for 10 minutes to prevent backflow. Immediately afterward, food was returned to the cage, and food intake was measured by weighing the remaining food at indicated time.

### Long-term intraperitoneal injections

To minimize the impact of stress responses on experimental outcomes, mice were acclimated via daily intraperitoneal injections one week prior to the study. During this habituation period, each mouse received daily intraperitoneal injections of saline (0.3 mL per injection). Following habituation, test compounds (LA-PTH or TIP-39) were administered intraperitoneally every other day. Injections were performed in the morning to minimize circadian rhythm effects. Food was provided immediately after each injection, and food intake was measured by weighing residual food at designated times. Body weight was monitored daily throughout this intraperitoneal injection period.

### Real-time PCR

Total RNA was extracted from hypothalamic tissue using Tissue Fast RNA Extraction Kit for Animal (ABclonal, #RK30120), and RNA concentration and purity were assessed with a NanoDrop 5500 spectrophotometer (Thermo Scientific). cDNA synthesis was performed using the PrimeScript RT Reagent Kit (ABclonal, #RK20433) according to the manufacturer’s instructions. Quantitative real-time PCR was carried out using the SYBR Green PCR system (Yeasen, #11184ES08) with the following thermal cycling conditions: 95°C for 2 minutes, followed by 40 cycles of 95°C for 10 sec and 60°C for 30 sec. Mouse β-actin was used as the endogenous control to normalize gene expression levels. Relative mRNA expression was calculated using the 2^−ΔΔCt^ method. Primer sequences were as follows:

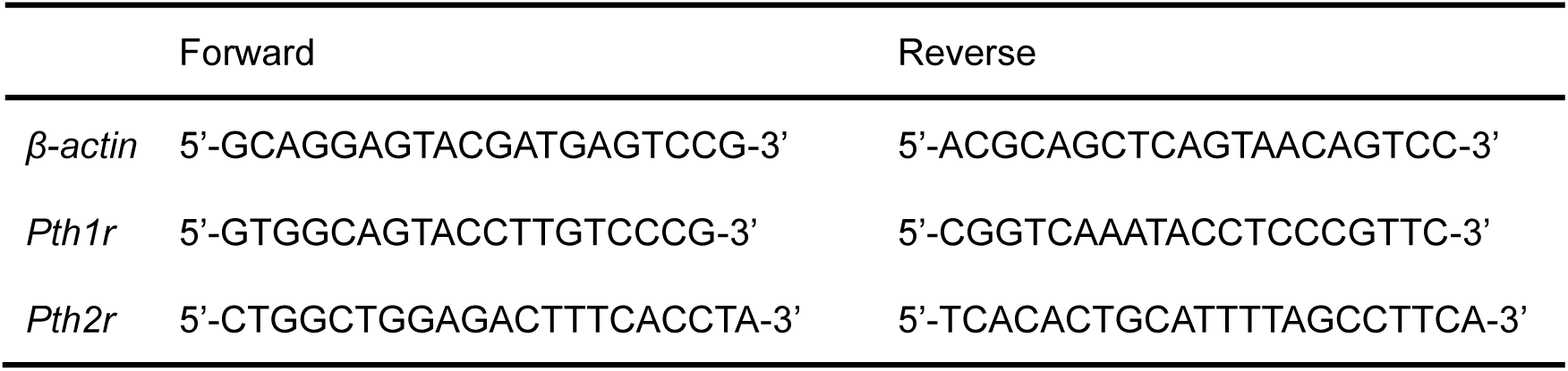

## OMICS DATA ANALYSIS AND BIOINFORMATICS

### Spectral library construction

We combined all DDA and DIA raw files acquired from tissues of four groups of mouse models to construct a FragPipe hybrid library using FragPipe (version 22.0) software. Data search was performed against a mouse Swiss-Prot database (Jul-2022, 17,114 entries) appended with iRT peptides and decoy sequences, and the unreviewed mouse olfactory and taste receptor sequences (1,328 entries). All runs were searched using MSFragger (version 4.1)^100^ with major settings: trypsin as the specific enzyme with two missed cleavages; C + 57.021464 as a fixed modification; M + 15.9949 and protein N-term + 42.0106 as variable modifications; precursor/fragment mass tolerances of 20 ppm; peptide length 7-50; maximum fragment charge of 2; minimum matched fragment ions 4. Post-search validation utilized Percolator and ProteinProphet, integrated within the Philosopher (version 5.1.1)^101^ software tool. 1% false discovery rate (FDR) thresholds were applied at peptide and protein levels.

Meanwhile, we also imported all raw files into Spectronaut (version 18.2) to generate a built-in hybrid library using the same mouse Swiss-Prot database with appended sequences. Trypsin and LysC were set as specific enzyme with maximum 2 missed cleavages, carbamidomethyl (C) was set to fixed modification, oxidation (M) and acetyl (protein N terminus) were set to variable modifications. 1% FDRs was set at peptide-spectrum match (PSM)/peptide/protein levels. Mass tolerances and iRT alignment were default settings.

### DIA MS data processing

For the analysis of MS data from normal mice, DIA raw files acquired from 22 tissues were processed using DIA-NN (version 1.8.1)^102^ in the library-free mode and Spectronaut (version 18.3)^103^ in the directDIA mode against the mouse Swiss-Prot database (Jul-2022, 17,114 entries), supplemented with the iRT peptide sequences, contaminant sequences, and the unreviewed olfactory and taste receptor sequences. The DIA-NN search parameters were set as default. Trypsin and LysC were set as specific enzyme with one missed cleavage, peptide length was set to 7 to 30, precursor charge state was 1 to 4 and precursor mass range was 300 to 1800 m/z. Quantification strategy was set to Robust LC (high precision), and cross-run normalization was RT-dependent. Precursor FDR was set at 1%. In DIA data search with Spectronaut, major default settings were applied: Trypsin and LysC were set as specific enzyme; Q value of precursor and protein cutoff 0.01; quantification based on MS2 area; normalization strategy set to automatic and mass tolerance was set to default. The protein quantification reports were exported for further analysis. According to our previous study on data search with DIA-NN and Spectronaut^104^, protein identifications from two platforms were combined to enhance the proteome coverage. Protein quantification data were primarily derived from DIA-NN search results, and supplemented with quantification of proteins exclusively present in Spectronaut search results.

For the analysis of MS data from mouse models, all DIA files acquired from four experimental groups were processed using either DIA-NN (version 1.8.1) with the FragPipe hybrid library or Spectronaut (version 18.2)^103^ with the built-in hybrid library. For data search with DIA-NN, the main search parameters were the same as the library-free mode except that the spectral library was imported separately. For data search with Spectronaut, the settings were described as above and consistent with the directDIA search configuration. Data search results with two platforms were combined in the same manner as described above.

### RNA-sequencing data analysis

After library quality control, quantified libraries were pooled and sequenced by using the Illumina comprehensive next-generation sequencing (NGS) technique, according to the effective library concentration and data amount. RNA sequencing libraries were prepared from total RNA samples using Trimmomatic (version 0.39) for adapter trimming and quality control. Raw paired-end reads were trimmed to remove adapters and low-quality bases, with a minimum length cutoff of 36 bp. Cleaned paired-end reads were aligned to the mouse reference genome (GRCm39) using the STAR (version 2.7.10) aligner. STAR was used for mapping the reads to the reference genome, ensuring accurate alignment to both the exonic and transcriptomic regions. Gene expression was quantified using featureCounts,^105^ and only reads mapping to the exonic regions were counted for different genes. Expression values were normalized using the Transcripts per Million (TPM) method.

### Curation of CMPs and protein/peptide ligands for GPCRs and RTKs

Cell membrane proteins (CMPs) were compiled from multiple curated sources. Reviewed UniProt entries annotated with the cell membrane (SL-0039) subcellular location were retrieved (3,187 entries). An addition of 237 reviewed catalytic receptors were retrieved from the IUPHAR/BPS Guide to PHARMACOLOGY. Moreover, 1,328 mouse olfactory and taste receptors were included. The final CMP inventory comprises 4,579 unique proteins.

For the re-analysis of DEPs and DEGs from previous studies^8, 9^, CMPs in different species were retrieved according to the UniProt annotation, supplemented with olfactory receptors of the same species, resulting in an inventory of 3,675 CMPs in *M. fascicularis* and 5,440 CMPs in *R. norvegicus*.

Protein or peptide ligands for GPCRs were curated from the ‘Physiological ligands’ category in GPCRdb^106^, and supplemented with endogenous ligands retrieved from the IUPHAR/BPS Guide to PHARMACOLOGY and Uniprot. Protein ligands for RTKs were also retrieved from the IUPHAR/BPS Guide and UniProt.

### Differential protein and gene analysis

Differential CMP analysis was performed using the limma package (v4.2) in R, which applies linear modeling and empirical Bayes moderation to improve variance estimation across replicates. Three pairwise comparisons were conducted between experimental groups (*i.e.* Chow vs Obs, Obs vs Exe, and T2D vs Obs). CMPs were considered differentially expressed proteins (DEPs) if they met either of the following criteria: (1) an absolute log2 fold change (|log_2_FC|) greater than 0.32 (*i.e.* FC > 1.25 in either direction), together with a Benjamini–Hochberg (BH) adjusted *p* < 0.05, or (2) detected in at least two replicates of one group but undetected in all replicates of the other group. We applied a modest FC cut-off similar to previous studies^107, 108^ in order to identify DEPs undergoing mild regulation to expand our candidate list for functional regulator discovery. Furthermore, reverse-trend DEPs are defined as CMPs or their protein ligands that are differentially expressed in one direction in the obesity vs chow comparison and in the opposite direction in the exercise vs obesity comparison.

Differential gene expression analysis was performed with the raw count data without prior normalization. Only genes with raw counts exceeding 10 in at least 30% of the samples were included in the analysis. Differential expression was conducted using the DESeq2 package in R. In the pairwise comparisons, similar to the DEP selection, genes with a |log_2_FC| > 0.58 (equivalent to FC > 1.5 in either direction) and a BH-adjusted *p* < 0.05 or genes exclusively detected in at least two replicates of one group and undetected in all replicates of the other group were considered as DEGs.

Overlapping of DEPs and DEGs reported in the same study was conducted based on their Ensembl gene IDs. Only proteins and genes that exhibit the same direction of change (up- or down-regulated) in the corresponding sample groups were considered overlapping DEP/DEG pairs. The fraction of DEPs overlapping with DEGs in the same tissue refers to the percentage of the number of overlapping DEP/DEG pairs over the total number of DEPs found in that tissue.

For the re-analysis of DEGs and DEPs reported from previous multi-omics studies^8, 9^, we filtered the original differential analysis reports to select DEPs and DEGs with the same FC and adjusted *p* cut-offs applied to our study. In the study of monkey models for obesity and T2D^8^, we re-analyzed DEGs and DEPs for the obesity group vs control group. In the study of a rat exercise model^9^, we re-analyzed DEGs and DEPs for the 8-week exercise group vs control group. All DEGs and DEPs were selected according to the CMP curation as described above.

### Principal component analysis (PCA)

PCA involves missing value imputation using the R package missMDA (version 1.19) and then dimensionality reduction using the R package FactoMineR (version 2.11). PCA was applied to the complete CMP proteome dataset, and the first and second principal components were extracted for further analysis. To evaluate group-wise differences, pairwise t-tests were conducted on the first and second principal components to detect significant shifts of CMP profiles in specific tissues between two experimental groups.

### GO and pathway enrichment

Biological Process analysis was performed for the top five tissues with the highest number of enriched CMPs using Gene Ontology Biological Process (GO-BP). For CMPs with low correlation between protein expression and mRNA expression (Spearman correlation < 0.3), GO enrichment analysis was also performed, focusing on Molecular Function (GO-MF). The analysis was carried out using the R package clusterProfiler, with pathways exhibiting a BH-adjusted *p* < 0.05 considered significantly enriched. Fold enrichment was calculated by dividing the GeneRatio (the number of genes associated with a specific pathway) by the BgRatio (the total number of genes in the background for that pathway). Pathway enrichment analysis based on differentially expressed CMPs was conducted for each tissue in three pair-wise comparisons. Reactome pathway enrichment was performed using the R package ReactomePA. Pathways with a BH-adjusted *p* < 0.01 were considered significantly enriched.

### HypoMap-based expression profiling of Pth1r and Pth2r

Single-cell RNA-seq datasets were obtained from the hypothalamic nucSeq resources hosted on CELLxGENE (available via https://www.mrl.ims.cam.ac.uk) and the integrated HypoMap Seurat object deposited in the Cambridge Apollo Repository (https://doi.org/10.17863/CAM.87955). All analyses were performed using Seurat (version 5.3.0) in R (version 4.5.1). Gene-positive cells were defined by a threshold of ≥1 UMI. UMAP embeddings were generated to resolve the distribution of Pth1r and Pth2r across annotated hypothalamic cell types and to quantify the proportion of gene-positive cells classified as neurons. Within receptor-positive neuronal subsets, co-expression with key hypothalamic markers (*Bdnf*, *Pomc*, *Agrp*, *Lepr*, *Glp1r*, *Mc3r*, *Crabp1*, *Mc4r* and the reciprocal *Pth1r*/*Pth2r*) were quantified as the fraction of double-positive cells relative to the corresponding receptor-positive neuronal populations.

### Statistical tests

Body weight measurements were analyzed using GraphPad Prism (GraphPad Software) with a two-sided unpaired t-test or two-way ANOVA. Blood glucose measurements were analyzed using a two-sided unpaired t-test. Food intake data were analyzed using two-way ANOVA to assess the effects of treatment and time. In metabolic cage experiments, heat production curves before and after 4°C cold exposure was used to calculate the area under curve (AUC), and AUC values were compared using two-way ANOVA. Statistical significance thresholds were defined as \**p* < 0.05, \*\**p* < 0.01, \*\*\**p* < 0.001, and \*\*\*\**p* < 0.0001.

## Notes

### Competing Interest Statement

The authors have declared no competing interest.

## References

1. McGee SL, Hargreaves M. Exercise adaptations: molecular mechanisms and potential targets for therapeutic benefit. Nat Rev Endocrinol 16, 495–505 (2020).

2. Lingvay I, Cohen RV, Roux CWl, Sumithran P. Obesity in adults. The Lancet 404, 972–987 (2024).

3. Abel ED, et al. Diabetes mellitus-Progress and opportunities in the evolving epidemic. Cell 187, 3789–3820 (2024).

4. Contrepois K, et al. Molecular Choreography of Acute Exercise. Cell 181, 1112–1130.e1116 (2020).

5. Emont MP, et al. A single-cell atlas of human and mouse white adipose tissue. Nature 603, 926–933 (2022).

6. Stocks B, et al. Integrated Liver and Plasma Proteomics in Obese Mice Reveals Complex Metabolic Regulation. Mol Cell Proteomics 21, 100207 (2022).

7. Sato S, et al. Atlas of exercise metabolism reveals time-dependent signatures of metabolic homeostasis. Cell Metab 34, 329–345.e328 (2022).

8. Zhang X, et al. A transcriptomic and proteomic atlas of obesity and type 2 diabetes in cynomolgus monkeys. Cell Reports 42, 112952 (2023).

9. Amar D, et al. Temporal dynamics of the multi-omic response to endurance exercise training. Nature 629, 174–183 (2024).

10. Amar D, et al. The mitochondrial multi-omic response to exercise training across rat tissues. Cell Metabolism 36, 1411–1429.e1410 (2024).

11. Aebersold R, Mann M. Mass-spectrometric exploration of proteome structure and function. Nature 537, 347–355 (2016).

12. Guo T, Steen JA, Mann M. Mass-spectrometry-based proteomics: from single cells to clinical applications. Nature 638, 901–911 (2025).

13. Santos R, et al. A comprehensive map of molecular drug targets. Nat Rev Drug Discov 16, 19–34 (2017).

14. Wang D, et al. A deep proteome and transcriptome abundance atlas of 29 healthy human tissues. Mol Syst Biol 15, e8503 (2019).

15. Li S, et al. Multiregional profiling of the brain transmembrane proteome uncovers novel regulators of depression. Science Advances 7, eabf0634 (2021).

16. Giansanti P, et al. Mass spectrometry-based draft of the mouse proteome. Nat Methods 19, 803–811 (2022).

17. Lu T, et al. Protein restriction reprograms the multi-organ proteomic landscape of mouse aging. Cell 188, 7309–7326.e7320. (2025).

18. Deng Y-T, et al. Atlas of the plasma proteome in health and disease in 53,026 adults. Cell 188, 253–271.e257 (2025).

19. Zhou Y, et al. TTD: Therapeutic Target Database describing target druggability information. Nucleic Acids Research 52, D1465–D1477 (2024).

20. Vogel C, Marcotte EM. Insights into the regulation of protein abundance from proteomic and transcriptomic analyses. Nature Reviews Genetics 13, 227–232 (2012).

21. Li W, et al. Turnover atlas of proteome and phosphoproteome across mouse tissues and brain regions. Cell 188, 2267–2287.e2221 (2025).

22. Davenport AP, Scully CCG, de Graaf C, Brown AJH, Maguire JJ. Advances in therapeutic peptides targeting G protein-coupled receptors. Nature Reviews Drug Discovery 19, 389–413 (2020).

23. Lorente JS, Sokolov AV, Ferguson G, Schiöth HB, Hauser AS, Gloriam DE. GPCR drug discovery: new agents, targets and indications. Nature Reviews Drug Discovery 24, 458–479 (2025).

24. Gottesman-Katz L, Latorre R, Vanner S, Schmidt BL, Bunnett NW. Targeting G protein-coupled receptors for the treatment of chronic pain in the digestive system. Gut 70, 970–981 (2021).

25. Chapman FA, Maguire JJ, Newby DE, Davenport AP, Dhaun N. Targeting the apelin system for the treatment of cardiovascular diseases. Cardiovasc Res 119, 2683–2696 (2023).

26. Ansari S, Khoo B, Tan T. Targeting the incretin system in obesity and type 2 diabetes mellitus. Nat Rev Endocrinol 20, 447–459 (2024).

27. Congreve M, de Graaf C, Swain NA, Tate CG. Impact of GPCR Structures on Drug Discovery. Cell 181, 81–91 (2020).

28. Waterson Michael J, Horvath Tamas L. Neuronal Regulation of Energy Homeostasis: Beyond the Hypothalamus and Feeding. Cell Metabolism 22, 962–970 (2015).

29. Ahrén B. Islet G protein-coupled receptors as potential targets for treatment of type 2 diabetes. Nat Rev Drug Discov 8, 369–385 (2009).

30. Fountas A, Diamantopoulos L-N, Tsatsoulis A. Tyrosine Kinase Inhibitors and Diabetes: A Novel Treatment Paradigm? Trends in Endocrinology & Metabolism 26, 643–656 (2015).

31. Barella LF, Jain S, Kimura T, Pydi SP. Metabolic roles of G protein-coupled receptor signaling in obesity and type 2 diabetes. FEBS J 288, 2622–2644 (2021).

32. Attwood MM, Fabbro D, Sokolov AV, Knapp S, Schiöth HB. Trends in kinase drug discovery: targets, indications and inhibitor design. Nature Reviews Drug Discovery 20, 839–861 (2021).

33. Pocai A. G protein-coupled receptors and obesity. Front Endocrinol (Lausanne) 14, 1301017 (2023).

34. Serdan TDA, et al. Impaired brown adipose tissue is differentially modulated in insulin-resistant obese wistar and type 2 diabetic Goto-Kakizaki rats. Biomedicine & Pharmacotherapy 142, 112019 (2021).

35. Castillo Í MP, Argilés JM, Rueda R, Ramírez M, Pedrosa JML. Skeletal muscle atrophy and dysfunction in obesity and type-2 diabetes mellitus: Myocellular mechanisms involved. Rev Endocr Metab Disord 26, 815–836 (2025).

36. Sakaue S, et al. A cross-population atlas of genetic associations for 220 human phenotypes. Nat Genet 53, 1415–1424 (2021).

37. Barton AR, Sherman MA, Mukamel RE, Loh PR. Whole-exome imputation within UK Biobank powers rare coding variant association and fine-mapping analyses. Nat Genet 53, 1260–1269 (2021).

38. Zhang Y, et al. Neuregulin4 Acts on Hypothalamic ErBb4 to Excite Oxytocin Neurons and Preserve Metabolic Homeostasis. Advanced Science 10, 2204824 (2023).

39. Grandl G, et al. Global, neuronal or β cell-specific deletion of inceptor improves glucose homeostasis in male mice with diet-induced obesity. Nature Metabolism 6, 448–457 (2024).

40. Cui H, López M, Rahmouni K. The cellular and molecular bases of leptin and ghrelin resistance in obesity. Nature Reviews Endocrinology 13, 338–351 (2017).

41. Petersen MC, Shulman GI. Mechanisms of Insulin Action and Insulin Resistance. Physiological Reviews 98, 2133–2223 (2018).

42. Graaf C, et al. Glucagon-Like Peptide-1 and Its Class B G Protein-Coupled Receptors: A Long March to Therapeutic Successes. Pharmacol Rev 68, 954–1013 (2016).

43. Zierath JR, He L, Gumà A, Odegoard Wahlström E, Klip A, Wallberg-Henriksson H. Insulin action on glucose transport and plasma membrane GLUT4 content in skeletal muscle from patients with NIDDM. Diabetologia 39, 1180–1189 (1996).

44. Krook A, et al. Characterization of signal transduction and glucose transport in skeletal muscle from type 2 diabetic patients. Diabetes 49, 284–292 (2000).

45. Rulifson IC, et al. Wnt signaling regulates pancreatic β cell proliferation. Proceedings of the National Academy of Sciences 104, 6247–6252 (2007).

46. Konstantinova I, et al. EphA-Ephrin-A-mediated beta cell communication regulates insulin secretion from pancreatic islets. Cell 129, 359–370 (2007).

47. Mellado-Gil J, et al. Disruption of Hepatocyte Growth Factor/c-Met Signaling Enhances Pancreatic β-Cell Death and Accelerates the Onset of Diabetes. Diabetes 60, 525–536 (2011).

48. Chen J, Zeng F, Forrester SJ, Eguchi S, Zhang M-Z, Harris RC. Expression and Function of the Epidermal Growth Factor Receptor in Physiology and Disease. Physiological Reviews 96, 1025–1069 (2016).

49. Goldberg IJ, et al. Decreased lipoprotein clearance is responsible for increased cholesterol in LDL receptor knockout mice with streptozotocin-induced diabetes. Diabetes 57, 1674–1682 (2008).

50. Jain S, Jacobson KA. Purinergic signaling in diabetes and metabolism. Biochemical Pharmacology 187, 114393 (2021).

51. Liu H, et al. Wnt Signaling Regulates Hepatic Metabolism. Science Signaling 4, ra6 (2011).

52. Hu L, Shan Z, Wang F, Gao X, Tong Y. Vascular endothelial growth factor B exerts lipid-lowering effect by activating AMPK via VEGFR1. Life Sciences 276, 119401 (2021).

53. Hilton C, Sabaratnam R, Drakesmith H, Karpe F. Iron, glucose and fat metabolism and obesity: an intertwined relationship. International Journal of Obesity 47, 554–563 (2023).

54. Miao R, Fang X, Zhang Y, Wei J, Zhang Y, Tian J. Iron metabolism and ferroptosis in type 2 diabetes mellitus and complications: mechanisms and therapeutic opportunities. Cell Death & Disease 14, 186 (2023).

55. Bugler-Lamb AR, et al. Adipocyte integrin-linked kinase plays a key role in the development of diet-induced adipose insulin resistance in male mice. Molecular Metabolism 49, 101197 (2021).

56. Lovászi M, Branco Haas C, Antonioli L, Pacher P, Haskó G. The role of P2Y receptors in regulating immunity and metabolism. Biochemical Pharmacology 187, 114419 (2021).

57. Wang Z, Wang QA, Liu Y, Jiang L. Energy metabolism in brown adipose tissue. The FEBS Journal 288, 3647–3662 (2021).

58. Green CD, Maceyka M, Cowart LA, Spiegel S. Sphingolipids in metabolic disease: The good, the bad, and the unknown. Cell Metab 33, 1293–1306 (2021).

59. Peirce V, Vidal-Puig A. Regulation of glucose homoeostasis by brown adipose tissue. Lancet Diabetes Endocrinol 1, 353–360 (2013).

60. Laplante MA, Monassier L, Freund M, Bousquet P, Gachet C. The purinergic P2Y1 receptor supports leptin secretion in adipose tissue. Endocrinology 151, 2060–2070 (2010).

61. Badeanlou L, Furlan-Freguia C, Yang G, Ruf W, Samad F. Tissue factor–protease-activated receptor 2 signaling promotes diet-induced obesity and adipose inflammation. Nature Medicine 17, 1490–1497 (2011).

62. Inoue E, et al. Beraprost sodium, a stable prostacyclin analogue, improves insulin resistance in high-fat diet-induced obese mice. J Endocrinol 213, 285–291 (2012).

63. Morigny P, Boucher J, Arner P, Langin D. Lipid and glucose metabolism in white adipocytes: pathways, dysfunction and therapeutics. Nature Reviews Endocrinology 17, 276–295 (2021).

64. Hagberg CE, Spalding KL. White adipocyte dysfunction and obesity-associated pathologies in humans. Nature Reviews Molecular Cell Biology 25, 270–289 (2024).

65. Wang B, et al. βAR-mTOR-lipin1 pathway mediates PKA-RIIβ deficiency-induced adipose browning. Theranostics 14, 5316–5335 (2024).

66. Pacher P, Bátkai S, Kunos G. The endocannabinoid system as an emerging target of pharmacotherapy. Pharmacol Rev 58, 389–462 (2006).

67. Rosenstock J, Hollander P, Chevalier S, Iranmanesh A. SERENADE: the Study Evaluating Rimonabant Efficacy in Drug-naive Diabetic Patients: effects of monotherapy with rimonabant, the first selective CB1 receptor antagonist, on glycemic control, body weight, and lipid profile in drug-naive type 2 diabetes. Diabetes Care 31, 2169–2176 (2008).

68. Shafiei-Jahani P, et al. CB2 stimulation of adipose resident ILC2s orchestrates immune balance and ameliorates type 2 diabetes mellitus. Cell Rep 43, 114434 (2024).

69. Li J, et al. Neurotensin is an anti-thermogenic peptide produced by lymphatic endothelial cells. Cell Metabolism 33, 1449–1465.e1446 (2021).

70. Ramirez-Virella J, Leinninger GM. The Role of Central Neurotensin in Regulating Feeding and Body Weight. Endocrinology 162, (2021).

71. Huang X, Yang C, Luo Y, Jin C, Wang F, McKeehan WL. FGFR4 Prevents Hyperlipidemia and Insulin Resistance but Underlies High-Fat Diet–Induced Fatty Liver. Diabetes 56, 2501–2510 (2007).

72. Zhen Y, et al. FGFR inhibition blocks NF-ĸB-dependent glucose metabolism and confers metabolic vulnerabilities in cholangiocarcinoma. Nat Commun 15, 3805 (2024).

73. Fafalios A, et al. A hepatocyte growth factor receptor (Met)-insulin receptor hybrid governs hepatic glucose metabolism. Nat Med 17, 1577–1584 (2011).

74. Bhushan B, Michalopoulos GK. Role of epidermal growth factor receptor in liver injury and lipid metabolism: Emerging new roles for an old receptor. Chem Biol Interact 324, 109090 (2020).

75. Luo X, et al. Reducing VEGFB expression regulates the balance of glucose and lipid metabolism in mice via VEGFR1. Mol Med Rep 26, 285 (2022).

76. Chui ZSW, Shen Q, Xu A. Current status and future perspectives of FGF21 analogues in clinical trials. Trends in Endocrinology & Metabolism 35, 371–384 (2024).

77. Jonker JW, et al. A PPARγ-FGF1 axis is required for adaptive adipose remodelling and metabolic homeostasis. Nature 485, 391–394 (2012).

78. Hagberg CE, et al. Targeting VEGF-B as a novel treatment for insulin resistance and type 2 diabetes. Nature 490, 426–430 (2012).

79. Livingstone C, Borai A. Insulin-like growth factor-II: its role in metabolic and endocrine disease. Clin Endocrinol (Oxf) 80, 773–781 (2014).

80. Seeley RJ, Drazen DL, Clegg DJ. The critical role of the melanocortin system in the control of energy balance. Annu Rev Nutr 24, 133–149 (2004).

81. Lawson EA. The effects of oxytocin on eating behaviour and metabolism in humans. Nature Reviews Endocrinology 13, 700–709 (2017).

82. Yoshimura M, Conway-Campbell B, Ueta Y. Arginine vasopressin: Direct and indirect action on metabolism. Peptides 142, 170555 (2021).

83. Liu T, Xu Y, Yi CX, Tong Q, Cai D. The hypothalamus for whole-body physiology: from metabolism to aging. Protein Cell 13, 394–421 (2022).

84. Steuernagel L, et al. HypoMap-a unified single-cell gene expression atlas of the murine hypothalamus. Nat Metab 4, 1402–1419 (2022).

85. Lei Y, et al. Region-specific transcriptomic responses to obesity and diabetes in macaque hypothalamus. Cell Metabolism 36, 438–453.e436 (2024).

86. Kobayashi K, et al. Endogenous ligand recognition and structural transition of a human PTH receptor. Mol Cell 82, 3468–3483.e3465 (2022).

87. Wang T, et al. Identification of a neural basis for energy expenditure in the mouse arcuate hypothalamus. Neuron 113, 3813–3833.e3819 (2025).

88. Wang X, et al. Molecular insights into differentiated ligand recognition of the human parathyroid hormone receptor 2. Proc Natl Acad Sci U S A 118, (2021).

89. Zhao LH, et al. Structure and dynamics of the active human parathyroid hormone receptor-1. Science 364, 148–153 (2019).

90. Feng X, et al. Parathyroid hormone alleviates non-alcoholic liver steatosis via activating the hepatic cAMP/PKA/CREB pathway. Front Endocrinol (Lausanne) 13, 899731 (2022).

91. Qin B, et al. Parathyroid hormone-related protein prevents high-fat-diet-induced obesity, hepatic steatosis and insulin resistance in mice. Endocr J 69, 55–65 (2022).

92. Jiang L, et al. A Quantitative Proteome Map of the Human Body. Cell 183, 269–283 e219 (2020).

93. Gardella TJ, Vilardaga JP. International Union of Basic and Clinical Pharmacology. XCIII. The parathyroid hormone receptors--family B G protein-coupled receptors. Pharmacol Rev 67, 310–337 (2015).

94. Dettori C, Ronca F, Scalese M, Saponaro F. Parathyroid Hormone (PTH)-Related Peptides Family: An Intriguing Role in the Central Nervous System. J Pers Med 13, 714 (2023).

95. Chen L, et al. Attenuation of Alzheimer’s brain pathology in 5XFAD mice by PTH1-34, a peptide of parathyroid hormone. Alzheimer’s Research & Therapy 15, 53 (2023).

96. Dobolyi A, Palkovits M, Usdin TB. The TIP39–PTH2 receptor system: Unique peptidergic cell groups in the brainstem and their interactions with central regulatory mechanisms. Progress in Neurobiology 90, 29–59 (2010).

97. Yavari M, Feizi A, Haghighatdoost F, Ghaffari A, Rezvanian H. The influence of parathyroidectomy on cardiometabolic risk factors in patients with primary hyperparathyroidism: a systematic review and meta-analysis. Endocrine 72, 72–85 (2021).

98. Izquierdo-Lahuerta A. The Parathyroid Hormone-Related Protein/Parathyroid Hormone 1 Receptor Axis in Adipose Tissue. Biomolecules 11, 1570 (2021).

99. MacLean B, et al. Skyline: an open source document editor for creating and analyzing targeted proteomics experiments. Bioinformatics 26, 966-968 (2010).

100. Kong AT, Leprevost FV, Avtonomov DM, Mellacheruvu D, Nesvizhskii AI. MSFragger: ultrafast and comprehensive peptide identification in mass spectrometry-based proteomics. Nat Methods 14, 513–520 (2017).

101. da Veiga Leprevost F, et al. Philosopher: a versatile toolkit for shotgun proteomics data analysis. Nat Methods 17, 869–870 (2020).

102. Demichev V, Messner CB, Vernardis SI, Lilley KS, Ralser M. DIA-NN: neural networks and interference correction enable deep proteome coverage in high throughput. Nat Methods 17, 41–44 (2020).

103. Bruderer R, et al. Extending the limits of quantitative proteome profiling with data-independent acquisition and application to acetaminophen-treated three-dimensional liver microtissues. Mol Cell Proteomics 14, 1400–1410 (2015).

104. Li S, et al. Generation of a Deep Mouse Brain Spectral Library for Transmembrane Proteome Profiling in Mental Disease Models. Molecular & Cellular Proteomics 23, 100777 (2024).

105. Liao Y, Smyth GK, Shi W. featureCounts: an efficient general purpose program for assigning sequence reads to genomic features. Bioinformatics 30, 923–930 (2014).

106. Herrera Luis PT, et al. GPCRdb in 2025: adding odorant receptors, data mapper, structure similarity search and models of physiological ligand complexes. Nucleic Acids Research 53, D425–D435 (2025).

107. Govaere O, et al. A proteo-transcriptomic map of non-alcoholic fatty liver disease signatures. Nat Metab 5, 572–578 (2023).

108. Needham EJ, et al. Personalized phosphoproteomics of skeletal muscle insulin resistance and exercise links MINDY1 to insulin action. Cell Metab 36, 2542–2559.e2546 (2024).

109. Chen T, et al. iProX in 2021: connecting proteomics data sharing with big data. Nucleic Acids Res 50, D1522–d1527 (2022).

110. Ma J, et al. iProX: an integrated proteome resource. Nucleic Acids Res 47, D1211–d1217 (2019).

111. Wang W, et al. The China National GeneBank Sequence Archive (CNSA) 2024 update. Hortic Res 12, uhaf036 (2025).

